# Plasmid manipulation of bacterial behaviour through translational regulatory crosstalk

**DOI:** 10.1101/2022.06.27.497698

**Authors:** Catriona M A Thompson, James P. J. Hall, Govind Chandra, Carlo Martins, Gerhard Saalbach, Susannah Bird, Samuel Ford, Richard H. Little, Ainelen Piazza, Ellie Harrison, Robert W. Jackson, Michael A. Brockhurst, Jacob G. Malone

**Affiliations:** Department of Molecular Microbiology, John Innes Centre, Colney Lane, Norwich, NR4 7UH, UK; School of Biological Sciences, University of East Anglia, Norwich Research Park, Norwich, Norfolk, NR4 7TJ, UK; Department of Evolution, Ecology and Behaviour Institute of Infection, Veterinary and Ecological Sciences University of Liverpool, Crown Street, Liverpool, L69 7ZB, UK; Department of Animal and Plant Sciences, University of Sheffield, Sheffield S10 2TN, UK; School of Biosciences, University of Birmingham, Edgbaston, Birmingham, B15 2TT, UK; Division of Evolution and Genomic Sciences, School of Biological Sciences, University of Manchester, Manchester M13 9PT, UK

**Author notes:** **Author Contributions**: MAB, JGM, RWJ, EH, JPJH obtained funding for the research. CMAT, JPJH, EH, RWJ, MAB and JGM designed the research. CMAT, JPJH, CM, GS, SB, SF, AP and RHL performed the research. JPJH and GC performed bioinformatic analyses. CMAT, MAB and SB analysed data. CMAT, JPJH, MAB and JGM wrote the paper.

**Keywords:** Translational regulation, *Pseudomonas*, Mobile Genetic Elements, Plasmid biology

## Abstract

Beyond their role in horizontal gene transfer, conjugative plasmids commonly encode homologues of bacterial regulators. Known plasmid regulator homologues have highly targeted effects upon the transcription of specific bacterial traits. Here, we characterise a plasmid translational regulator, RsmQ, capable of taking global regulatory control in *Pseudomonas fluorescens* and causing a behavioural switch from motile to sessile lifestyle. RsmQ acts as a global regulator controlling the host proteome through direct interaction with host mRNAs and interference with the host’s translational regulatory network. This mRNA interference leads to largescale proteomic changes in metabolic genes, key regulators and genes involved in chemotaxis, thus controlling bacterial metabolism and motility. Moreover, comparative analyses found RsmQ on a large number of divergent plasmids isolated from multiple bacterial host taxa, suggesting the widespread importance of RsmQ for manipulating bacterial behaviour across clinical, environmental, and agricultural niches. RsmQ is a widespread plasmid global translational regulator primarily evolved for host chromosomal control to manipulate bacterial behaviour and lifestyle.

**Significance Statement:** Plasmids are recognised for their important role in bacterial evolution as drivers of horizontal gene transfer. Less well understood are the effects of plasmids upon bacterial behaviours by manipulating the expression of key bacterial phenotypes. Until now, examples of plasmid manipulation of their bacterial hosts were limited to highly targeted transcriptional control of a few related traits. In contrast, here we describe the first plasmid global translational regulator evolved to control the bacterial behavioural switch from a motile to a sessile lifestyle and bacterial metabolism, mediated through manipulation of the bacterial proteome. Moreover, this global translational regulator is common across divergent plasmids in a wide range of bacterial host taxa, suggesting that plasmids may commonly control bacterial lifestyle in the clinic, agricultural fields, and beyond.

## Introduction

Bacteria regulate the expression of functional traits in response to their environment, enabling colonisation of diverse ecological niches. However, control over bacterial gene regulation is not exclusively under the control of the bacterial genome. The mobile genetic elements which inhabit bacterial hosts, such as conjugative plasmids, commonly encode homologues of bacterial regulators. The introduction of plasmid-encoded regulator homologues into the bacterial cell can rewire the gene regulatory networks of the bacterium, potentially altering the expression of bacterial traits; a process termed plasmid-chromosome crosstalk (PCC,). However, how and why plasmid-encoded regulators would manipulate the expression of bacterial traits is poorly understood.

To date, well-characterised PCCs involve plasmid-encoded transcriptional regulators that alter the expression of specific bacterial traits. For example, in *Acinetobacter baumannii* several multidrug resistance plasmids encode transcriptional repressors of the bacterial type VI secretion system (T6SS) (1). Plasmid-mediated repression of the T6SS enhances plasmid horizontal transmission by ensuring that plasmid recipient cells are not killed by the donor’s T6SS apparatus (2). Similarly, plasmid encoded transcriptional regulators alter the expression of several chromosomal regulators of virulence associated traits in *Rhodococcus equi*, thus enhancing survival of both the bacterium and the plasmid in macrophages by stalling phagosomal maturation (3). Together these examples suggest that plasmid encoded regulatory homologues may have important fitness consequences for the plasmid, either through horizontal replication, through conjugation to new host cells or through vertical replication within the current host cell (4).

The molecular mechanisms of known PCC involve plasmid-encoded transcriptional regulators causing targeted changes to the expression of small numbers of chromosomal genes. Although transcriptional regulation is important for bacterial survival and adaptation, bacteria also rely on translational regulation to respond to changes in their environment (5). Bacteria are able to exert this control by deploying second messenger signals (6), directly altering the ribosome (7) or impacting mRNA stability and accessibility via pathways such as Gac-Rsm (8, 9). It is currently unknown whether conjugative plasmids are able to manipulate translational regulatory pathways. The Gac-Rsm pathway is one of the best characterised translational regulatory systems in pseudomonads (10–14) and controls a wide variety of traits including biofilm formation (15), motility (16), quorum sensing (17), siderophore production (18) and virulence (8, 19). Gac-Rsm is highly conserved within the *Pseudomonas* genus and comparable systems exist in a wide range of bacteria (8, 14, 18, 20, 21). Rsms are small (9 kDa) proteins that are able to interact directly with the bases AnGGA around the ribosome binding site (RBS) of their target transcript (21–24). Rsm proteins can both activate and repress bound mRNA transcripts, either by opening up the mRNA to allow ribosomal access to the RBS, or by making the RBS inaccessible (24–26). This allows Rsm proteins to exert tight translational control over a wide range of targets to impact bacterial phenotypes (27). The activation of Rsm is regulated by the activation of the GacA/S two-component system (TCS) which is activated by a complex but largely uncharacterised set of environmental cues. Upon activation, GacA promotes transcription of the small-regulatory RNAs RsmY and RsmZ, which leads to the sequestering of regulatory Rsm proteins away from their mRNA targets through competition for binding (24, 28). The number of Rsm proteins encoded by individual *Pseudomonas* species varies, with each protein having both unique and overlapping regulons with other Rsm proteins (29). The large number of traits regulated by the Gac-Rsm system suggests that there could be significant effects on bacterial behaviour caused by PCC manipulating this system.

In this study, we investigate the role of translational regulation in mediating PCC between *Pseudomonas fluorescens* SBW25 and the 425 kb conjugative plasmid pQBR103. *P. fluorescens* is a common, soil-dwelling, plant growth promoting bacterium that is capable of accepting diverse plasmids, including those from the pQBR family of large conjugative plasmids (30, 31). Both SBW25 and the pQBR plasmids were first isolated in the 1990s from the sugar beet rhizosphere at Wytham Woods in the United Kingdom (30, 32) . The ability of several of the pQBR plasmids to persist within *P. fluorescens* strains across a range of environments including in soil, on plants, and in lab media has been well documented (30, 31, 33, 34). Moreover, acquisition of pQBR103 by *P. fluorescens* SBW25 alters the expression of ∼440 chromosomal genes (34, 35). The large-scale regulatory disruption caused by pQBR103 can be negated by a range of compensatory mutations restoring wild-type (WT) expression levels, including loss of function mutations affecting the bacterial TCS *gacA/S*. Notably, while the genetic sequence of pQBR103 encodes a range of accessory functions including mercury resistance and UV resistance, it also encodes a homologue of the widespread *rsmA* bacterial translational regulator gene, which we identify here as *rsmQ*.

To understand the function of *rsmQ*, we explored the distribution of *rsm* genes on plasmids, and the effects of *rsmQ* on the transcriptome and proteome of SBW25, as well as on the expression of key bacterial ecological traits. Further, we biochemically characterised the interactions of RsmQ with a close structural proxy for RNA (single stranded DNA (ssDNA)) and with the bacterial Rsm proteins. Our findings show that *rsm* genes are widespread on *Pseudomonas* plasmids, and that RsmQ interacts with the resident Gac-Rsm system and the host RNA pool, binding to specific nucleotide motifs in order to post-transcriptionally regulate translation. RsmQ extensively remodelled the SBW25 proteome including altering production of metabolism, nutrient uptake and chemotaxis pathways, despite having only a limited impact on the SBW25 transcriptome. In turn, RsmQ translational regulation altered the expression of ecologically important bacterial behaviours, most notably motility and carbon source metabolism. Together our findings expand the known molecular mechanisms causing plasmid-chromosome crosstalk to include translational regulator homologues, which act in this case to extensively manipulate bacterial behaviour by altering the expression of a range of ecologically important bacterial traits. These findings have broad implications for understanding the role of plasmids in microbial communities.

## Results

### Plasmids encode regulatory protein homologues

The ORF *PQBR443*, hereafter *rsmQ*, was identified on pQBR103 as a homologue of the chromosomal *csrA/rsmA* genes found widely within proteobacteria. We hypothesised that this gene could act as a mediator of PCC (36). To identify whether carriage of an *rsm* homologue is peculiar to pQBR103 or is a general phenomenon across plasmids, we investigated the distribution of *rsmQ* homologues in the 12,084 plasmids of the COMPASS database (37). Within this set, and consistent with previous studies (38), we detected 106 putative *rsmQ* homologues on 98 plasmids (0.8%), mostly isolated (92/98) from proteobacteria, particularly Pseudomonadaceae and Legionellaceae (Figure 1a). The distribution of *rsm*-containing plasmids was not uniform across taxa (Fisher’s Exact Test, p < 0.0005): approximately 20% of Pseudomonadaceae (41/196) and Piscirickettsiaceae (14/72) plasmids, and over 50% of Legionellaceae (18/30) plasmids contained *rsm* homologues, while no *rsm* homologues were detected on any of the 3621 Enterobacteriaceae plasmids. *rsm*-containing plasmids were relatively large, with the smallest at 32.4 kb, sitting at the larger end of the size distribution for each taxon (Figure S1a). There was no general association between *rsm*-carriage and plasmid mobility across taxa, although within Legionellaceae, *rsm*-encoding plasmids tended to be conjugative (Fisher’s Exact Test Bonferroni-adjusted p-value = 0.02). Within Pseudomonadaceae, *rsm-*encoding plasmids tended to have proportionally more genes with predicted *rsm* binding sites (Kolmogorov-Smirnov test, p = 0.012, Figure S1b). Overall, these patterns suggest that plasmid carriage of *rsm* is not uncommon, but is taxon-specific, indicating a functional role that is associated with particular groups of microorganisms.

**Figure 1.**
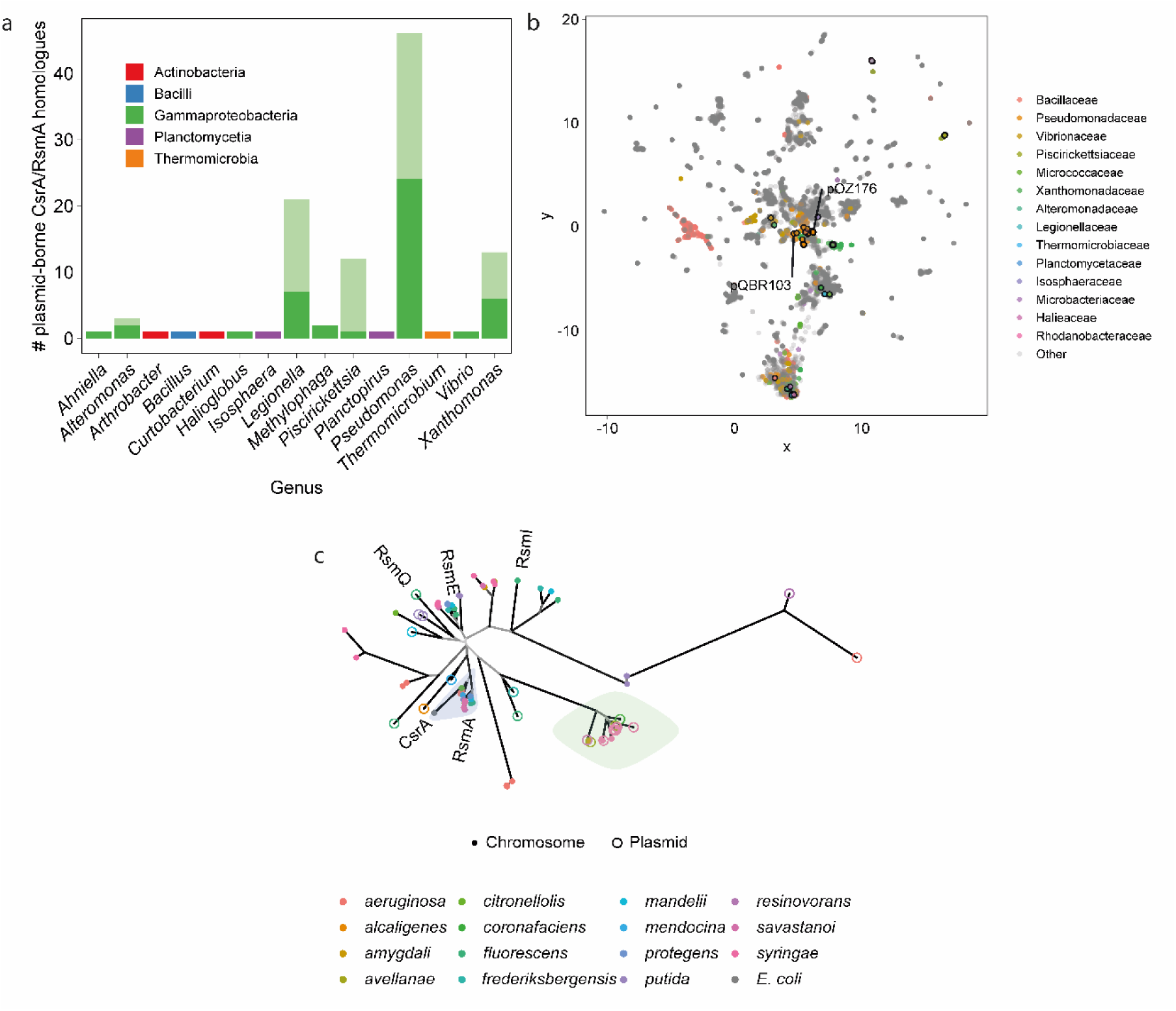
RsmQ is found on a wide range of conjugative plasmids. a) Taxonomic distribution of plasmid borne *csrA/rsmA* homologues identified in COMPASS. The paler part of each stacked bar indicates genes that were identical at the nucleotide level to other identified homologues. b) COMPASS plasmid diversity represented by non-metric multidimensional scaling of MASH sequence distances. Families with ≥1 plasmid with a *csrA/rsmA* homologue are coloured according to the legend. Plasmids encoding *csrA/rsmA* homologues are outlined in black. Selected plasmids from various taxa are annotated. c) Unrooted phylogenetic tree of *Pseudomonas csrA/rsmA* homologues from COMPASS, with corresponding chromosomal homologues (where available) and genes from selected reference strains. Branches leading to nodes with >80% bootstrap support are coloured black, with decreasing support indicated with increasingly pale grey. *P. fluorescens* SBW25 *csrA/rsmA* homologues are labelled, as is pQBR103 *rsmQ*, and *E. coli csrA*. The blue highlight indicates a well-supported (bootstrap support 0.84) group of *rsmA*-like homologues. The green highlight indicates the group of related plasmid and chromosomal genes from plant pathogen *Pseudomonas* discussed in the text.

It is possible that *rsm*-encoding plasmids have recently spread horizontally across different species. If this was the case, we would expect *rsm*-encoding plasmids to be more similar to one another than to non-*rsm*-encoding plasmids within each taxon. To investigate the diversity of *rsm*-encoding plasmids relative to the other plasmids in COMPASS, we performed UMAP non-metric multidimensional scaling (NMDS) on pairwise MASH distances between plasmids (39–41). Within the diversity of plasmids in COMPASS, *rsm*-containing plasmids are diverse and often cluster close to non-*rsm*-containing plasmids isolated from the same taxa (Figure 1b), suggesting that carriage of *rsm* regulators by plasmids is a convergent trait that has emerged several times over.

Global regulatory genes may be frequently (re)acquired by plasmids from the bacterial chromosome. Alternatively, these genes may have a prolonged association with plasmids and evolve distinctly to chromosomal genes. To investigate these possibilities, we built a phylogenetic tree of the *csrA/rsmA* homologues from all *Pseudomonas* plasmids and their associated chromosomes (where available), alongside the *rsm* genes from seven diverse *Pseudomonas* strains: *Pseudomonas protegens* CHA0, *P. fluorescens* Pf0-1, *P. protegens* Pf-5, *P. fluorescens* SBW25, *Pseudomonas putida* KT2440, *Pseudomonas aeruginosa* PAO1, and *P. aeruginosa* PA14 (Figure 1c). Chromosomal homologues of *csrA/rsmA* formed several distinct clusters (bootstrap support >80%), with one cluster including *P. fluorescens* SBW25 *rsmA* and the *E. coli* homologue *csrA*. However, plasmid-borne *csrA/rsmA* homologues were more divergent than those that were chromosomally encoded (Figure 1c). Additionally, chromosomal homologues (including the *P. fluorescens* SBW25 genes *rsmE* and *rsmI*) formed a distinct cluster. Consistent with the phylogenetic analysis, sequence variation among chromosomal *rsm* homologues was significantly lower than when comparing chromosomal-with plasmid-borne *rsm* homologues (Wilcoxon test, Bonferroni-adjusted p < 0.0001), or when comparing plasmid-borne *rsm* homologues with one another (Wilcoxon test, p < 0.0001). The principal exception to this pattern was a cluster of closely related plasmid and chromosomal *rsm* genes from plant pathogenic *Pseudomonas* (green highlighted, figure 1c). However, it is possible that some of these chromosomal variants are associated with chromosomally located mobile genetic elements, as at least one of these homologues is located on an annotated integrative conjugative element (42).

Overall, our comparative sequence analysis suggests that diverse plasmids have independently acquired *rsm* homologues, which then evolve and diversify as part of the plasmid mobile gene pool, distinct from their chromosomal counterparts. Although plasmid-encoded *rsm* homologues are widespread among plasmids (38), very little is currently known about their role in PCC or how they might impact bacterial behaviour.

### RsmQ binds to specific RNA targets

Despite a high degree of sequence similarity, it was unknown if RsmQ would be functionally similar to the chromosomally encoded SBW25 Rsm proteins (RsmA/E/I). Rsm proteins from *Pseudomonas* species interact with a conserved RNA sequence (AnGGA), with these bases interacting directly with the proteins’ conserved binding site (VHRE/D) (23, 24). To confirm whether RsmQ acts similarly, we designed a high throughput method to examine the nucleotide binding properties of RsmQ *in vitro* using the ReDCaT surface plasmon resonance (SPR) system (43), which is primarily designed for examining dsDNA interactions. Because Rsm proteins only interact with the nucleotide bases of RNA molecules, protein-nucleic acid interactions can be effectively examined using ssDNA probes. ssDNA probes containing the predicted RNA target sequence (ACGGA) and a non-specific scrambled sequence were synthesised with the ReDCaT linker at the 3’ end, with either a linear or a hairpin secondary structure with the potential binding site at the top of the hairpin.

RsmQ interacted strongly with both the minimal (GGA) and full length (ACGGA) binding sites when these were at the top of a hairpin loop. When the binding sequence was presented in a linear oligo RsmQ could interact but quickly dissociated, suggesting that the preferred binding site is open at the top of a hairpin loop (Figure S2). No interaction was seen between RsmQ and a scrambled binding site confirming that the binding is specific to the target GGA/ACGGA sequence (Figure 2).

**Figure 2:**
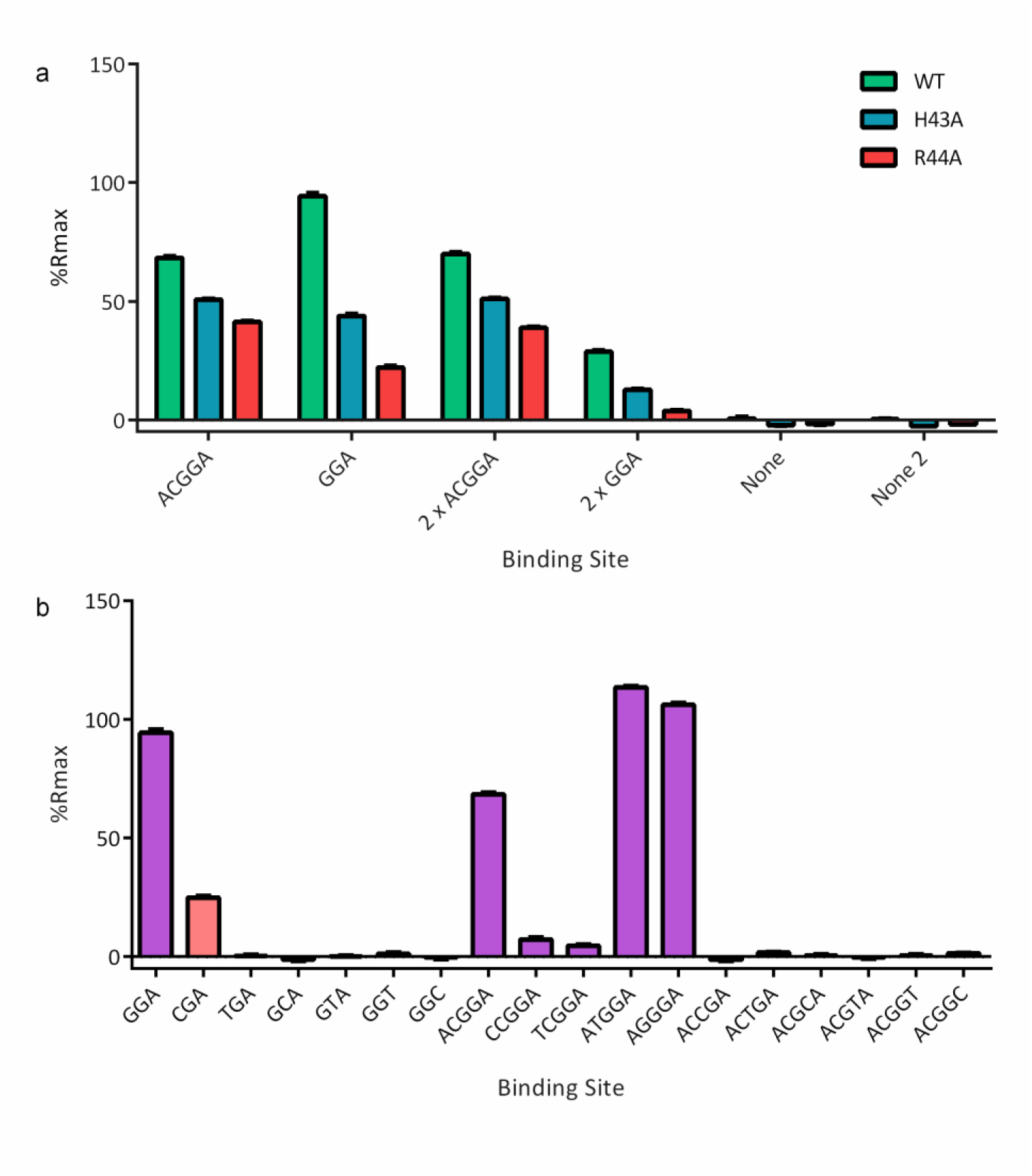
RsmQ interacts with its preferred binding sequence (GGA/AnGGA) and this interaction is mediated by the VHRE/D binding site. a) Percentage R_max_ values for RsmQ WT (green) H43A (blue) and R44A (red) binding to ssDNA containing the binding sites shown above. b) Percentage R_max_ values for WT RsmQ binding to ssDNAs containing the above binding site. Binding sites that are predicted to bind RsmQ are shown in purple. All oligos are designed as hairpins and results shown are for RsmQ at a concentration of 100 nm.

Next, we tested if the RNA binding interaction was co-ordinated by the conserved VHRD/E motif at the C-terminus of RsmQ by examining the binding of two RsmQ mutants (H43A and R44A) to the ssDNA probes. The alteration of these residues significantly reduced the efficiency of RsmQ binding to the target sequence (Figure 2a). Finally, to confirm the minimum RNA binding sequence a series of near-identical ssDNA nucleotides were tested containing the simple and full binding site sequences with a single base pair change in each case. RsmQ preferentially bound to the known binding sites GGA and A(N)GGA with a markedly higher affinity than to any of the alternate sequences tested and with a slight preference for ATGGA/AGGGA sequences, further supporting the hypothesis that RsmQ is a specific RNA binding protein that functions similarly to the chromosomal Rsm proteins (Figure 2b).

### RsmQ post-transcriptionally regulates the abundance of metabolism, nutrient transport, and chemotaxis proteins

To examine the impact of RsmQ on SBW25 regulation, a plasmid lacking *rsmQ* was created by allelic exchange in a kanamycin resistance gene-containing variant of pQBR103 (pQBR103^Km^) and freshly conjugated into SBW25. Expression of the chromosomal *rsm* genes generally peaks as growing cells make the switch from exponential to stationary phase growth (44). Therefore, *P. fluorescens* SBW25 cells carrying pQBR103^Km^ or pQBR103^Km^-Δ*rsmQ* or no plasmid were grown to late exponential phase (OD_600_ = 1.4) in KB liquid medium and a combination of RNA-seq and TMT quantitative proteomics were used to examine changes in mRNA and protein abundances. Five independent transconjugants for each plasmid as well as five independent colonies of the plasmid-free strain were used for these experiments.

Contrary to previous studies of exponentially growing cultures where carriage of pQBR103 caused large-scale transcriptional changes (35, 36, 45), we observed only modest transcriptional changes associated with plasmid carriage under our growth conditions, with only 54 genes upregulated more than two-fold and 33 downregulated by pQBR103^Km^ carriage, and similar numbers seen for pQBR103^Km^ -Δ*rsmQ*. Genes whose transcription was upregulated predominantly encoded transport-related membrane proteins, such as ABC and glutamine high-affinity transporter components. In addition, *PFLU1887* and *PFLU1888* encoding components of a putative transposase, were upregulated by pQBR103^Km^ carriage, consistent with previous studies (46). Genes downregulated by plasmid acquisition included several cytochrome C oxidases and metabolic enzymes such as L-lactate and shikimate dehydrogenases. pQBR103^Km^ and pQBR103^Km^-Δ*rsmQ* had highly similar transcriptional effects, suggesting that that RsmQ had little impact on the transcription of either chromosomal or plasmid genes.

In contrast, we observed major differences between the proteomes of SBW25 (pQBR103^Km^) and SBW25 (pQBR103^Km^-Δ*rsmQ*), confirming that RsmQ is indeed a post-transcriptionally acting translational regulator. Specifically, 581 SBW25 proteins showed at least two-fold increased abundance in the absence of *rsmQ* (i.e. their translation is suppressed by RsmQ) and 203 showed at least two-fold decreased abundance (Figure 3a and Table S1). Intriguingly, RsmQ regulation predominantly affected the host proteome, with the abundances of only a small fraction of plasmid-encoded proteins (16/733) altered by the presence of *rsmQ*.

**Figure 3:**
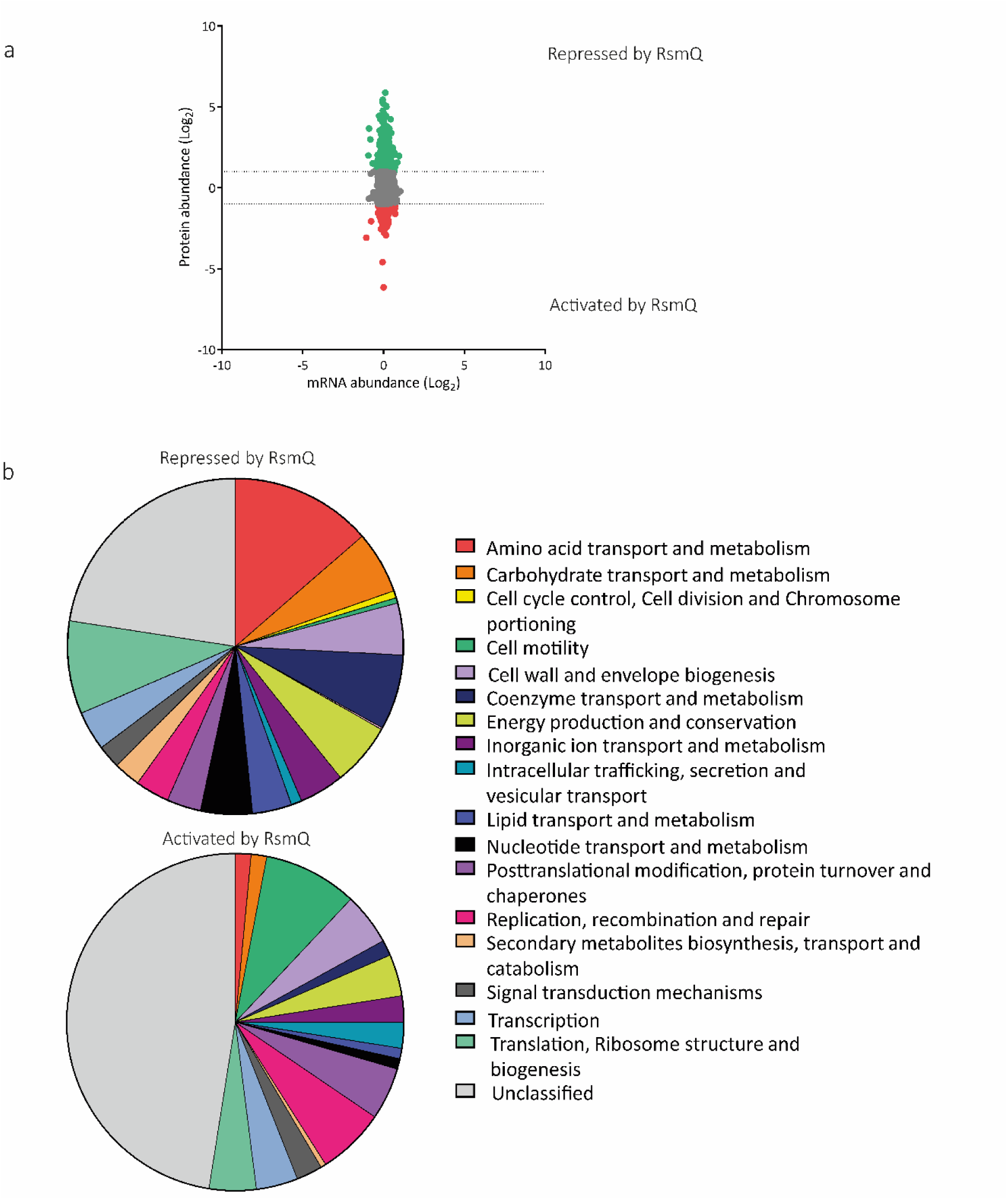
The loss of RsmQ causes widescale proteomic changes. a) Comparative scatter plot comparing log_2_-fold mRNA abundance changes from RNAseq (n=5) to protein abundance changes seen by TMT quantitative proteomics (n=3). b) COG categories of proteins that showed a greater than log_2_-fold change when *rsmQ* is lost (Repressed by RsmQ = 581, Activated by RsmQ = 203).

We next determined the COG functional categories of SBW25 proteins whose abundances were altered by the presence of RsmQ (Figure 3b). The 581 proteins downregulated by RsmQ were disproportionately associated with amino acid, coenzyme and carbohydrate transport and metabolism, as well as proteins involved in mRNA translation and ribosome stability. Among the 50 most strongly downregulated proteins we observed multiple inorganic ion transporters and receptor proteins, for example pyoverdine receptors and iron transporters (PFLU3378, PFLU2545 (FpvA), PFLU0295) (47–50), a copper transport outer-membrane porin (PFLU0595 (OprC homolog)) and the PhoU phosphate ABC transporter known to repress the Pho operon (PFLU6044 (51)), alongside proteins involved in amino-acid transport (e.g. ABC transporters PFLU0827 & PFLU0332 and metabolism (e.g. GlyA – PFLU565). Together these data suggest a role for RsmQ in the control of SBW25 nutrient acquisition and metabolism.

Conversely, proteins upregulated by RsmQ included a large number of motility proteins and DNA recombination and repair systems, alongside a larger fraction of uncharacterised proteins than in the RsmQ downregulated group of proteins. The most strongly upregulated proteins included a striking number of chemotaxis pathway components. In addition to CheA (PFLU4414) we identified 5 putative methyl-accepting chemotaxis proteins (e.g. PFLU2358, PFLU3427 & PFLU2486). RsmQ upregulated proteins also included the master-regulator of motility FleQ (PFLU4443 (52, 53)) and an uncharacterised RpiR family transcriptional regulator (PFLU257). This suggests that RsmQ modifies bacterial motility through altering cellular perception of the environment and the availability of local nutrient sources, as well as by directly controlling production of motility apparatus. Interestingly, the Gac-Rsm TCS repressor protein RetS (PFLU0610 (54)) is also upregulated by RsmQ, supporting a further regulatory linkage between pQBR103^Km^ carriage, RsmQ function and the Gac/Rsm pathway.

The global regulator Hfq (PFLU0520) was shown to be downregulated in a Δ*rsmQ* background, suggesting an additional level of post-transcriptional regulatory control. A fraction of the published SBW25 Hfq regulon (55) was shown to be up/down-regulated in the Δ*rsmQ* background. However, only relatively modest regulatory overlap was observed between the two systems. Intriguingly, one of the few plasmid-encoded proteins that was significantly affected by RsmQ was an Hfq homologue (pQBR0137), whose abundance increased upon *rsmQ* deletion (Table S1). Only the chromosomally encoded Hfq possesses a predicted RsmQ binding site however, suggesting that the plasmid-borne protein may be compensating for reduced chromosomal Hfq levels under conditions where RsmQ is not functional.

Sequence analysis suggests that around 50% of the genes encoding proteins whose abundance is differentially regulated by RsmQ contained an AnGGA binding site upstream, or within the first 100 bp of the ORF, with an additional 25% of all genes containing the simpler GGA motif. This pattern is consistent with RsmQ binding to these mRNAs to regulate their translation. To test this, we next designed ssDNA probes to examine if RsmQ indeed targeted the binding sites of the genes predicted to be directly regulated. Binding site probes were designed to be 30 bp long with the predicted binding site in the centre of the oligo with the ReDCaT linker on the 3’ end. As previously described SPR was performed to determine if an interaction occurred between the sequence and RsmQ *in vitro.* Using the hairpin AnGGA synthetic binding site oligos as a guide, five of the potential binding site oligos showed a %R_max_ of greater than 50% (Figure 4). These binding sites were the upstream regions of PFLU0923 (ATGGA), PFLU3378 (AGGGA), PFLU1516 (AGGGA), PFLU4443 (AGGGA, FleQ) and PFLU4726 (ATGGA) with the highest binding seen with PFLU3378.

**Figure 4:**
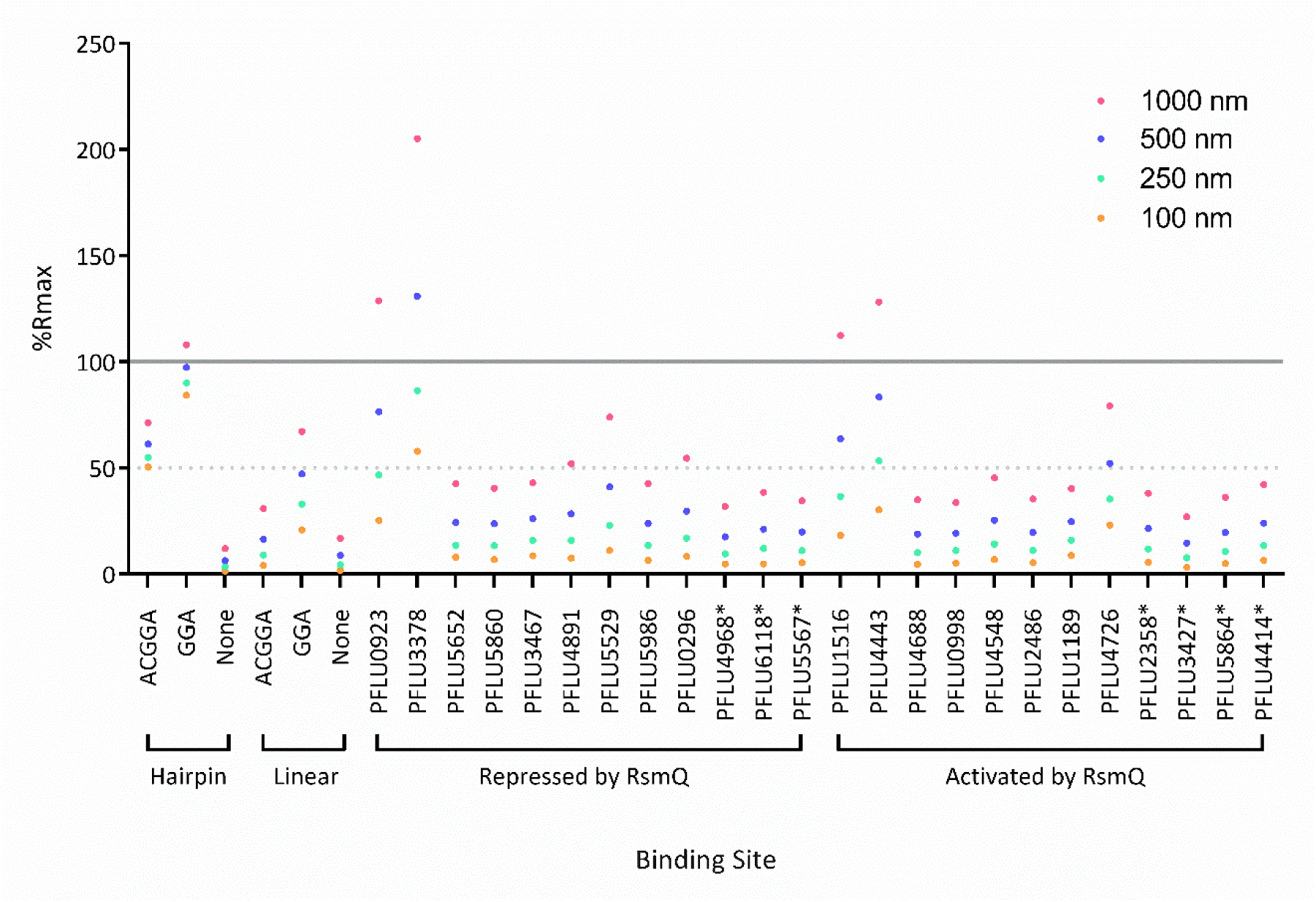
RsmQ binds to the upstream regions of predicted mRNA targets. Purified RsmQ was tested against ssDNA oligos with synthetic oligos run as a control. Oligos are labelled as in table S4, with the genetic identifiers used to indicate the binding sites associated with those ORFs. Genes annotated with a * indicate ORFs that contain a simplified GGA binding site but are not predicted to have a full Rsm (AnGGA) binding site within 500 bp upstream or the first 100 bp of the ORF. Percentage R_max_ values are shown for four concentrations. A solid line indicates 100% R_max_ and a dotted line for 50% R_max_. 100% R_max_ suggests 1:1 ligand protein binding as the experimentally acquired response is equal to the predicted response, with a 50% R_max_ suggesting a weaker interaction or a 2:1/1:2 response.

Despite an apparent preference for ATGGA/AGGGA binding sites, it is likely that the secondary structure was the overriding predictor of binding. Secondary structure predictions suggested that PFLU3378 was the only oligo with the binding site fully open at the top of the stem loop, with the rest showing partial occlusion of the ssDNA binding site by incorporation into a stem loop. These data confirm direct interaction between RsmQ and at least some of its predicted targets and further support the importance of mRNA secondary structure for successful RsmQ binding.

### RsmQ interacts with the host Rsm system

Notwithstanding the evidence for direct regulation of translation by RsmQ binding to mRNA, the large remainder of differentially regulated proteins without predicted Rsm binding sites suggests an indirect mechanism by which RsmQ regulates the abundance of these proteins. Given that RsmQ closely mimics the RNA binding characteristics of host Rsm proteins (Figure 2), we next investigated whether RsmQ interacts with other elements of the host Rsm regulatory pathway.

The activity of host Rsm proteins is controlled by the ncRNAs RsmY/Z, which act as protein sponges, sequestering Rsm proteins away from their target mRNAs. To test RsmQ binding to the ncRNAs RsmY and RsmZ, we copied the individual stem loops of each ncRNA into ssDNA oligos of approximately 25bp in length and attached them to the ReDCaT linker. The oligos were modelled to determine the location of the binding site in both the ncRNA and in the case of the ssDNA sequence, to determine if this was located at the top of a stem loop. Strong binding to several ssDNA probes was observed, in each case contingent on the presence of at least a GGA sequence at the top of a stem loop, with RsmY 1-25 and RsmZ 26-50 having AnGGA motifs present (Figure 5a). These data suggest that RsmQ interacts with the Gac-Rsm regulatory system by binding to the host ncRNAs RsmZ and RsmY. This would lead to either an increase in RsmQ target translation as RsmQ is titrated away from its targets, or an increase in RsmA/I/E binding to target mRNAs due to a reduction in available RsmZ/Y binding sites.

**Figure 5:**
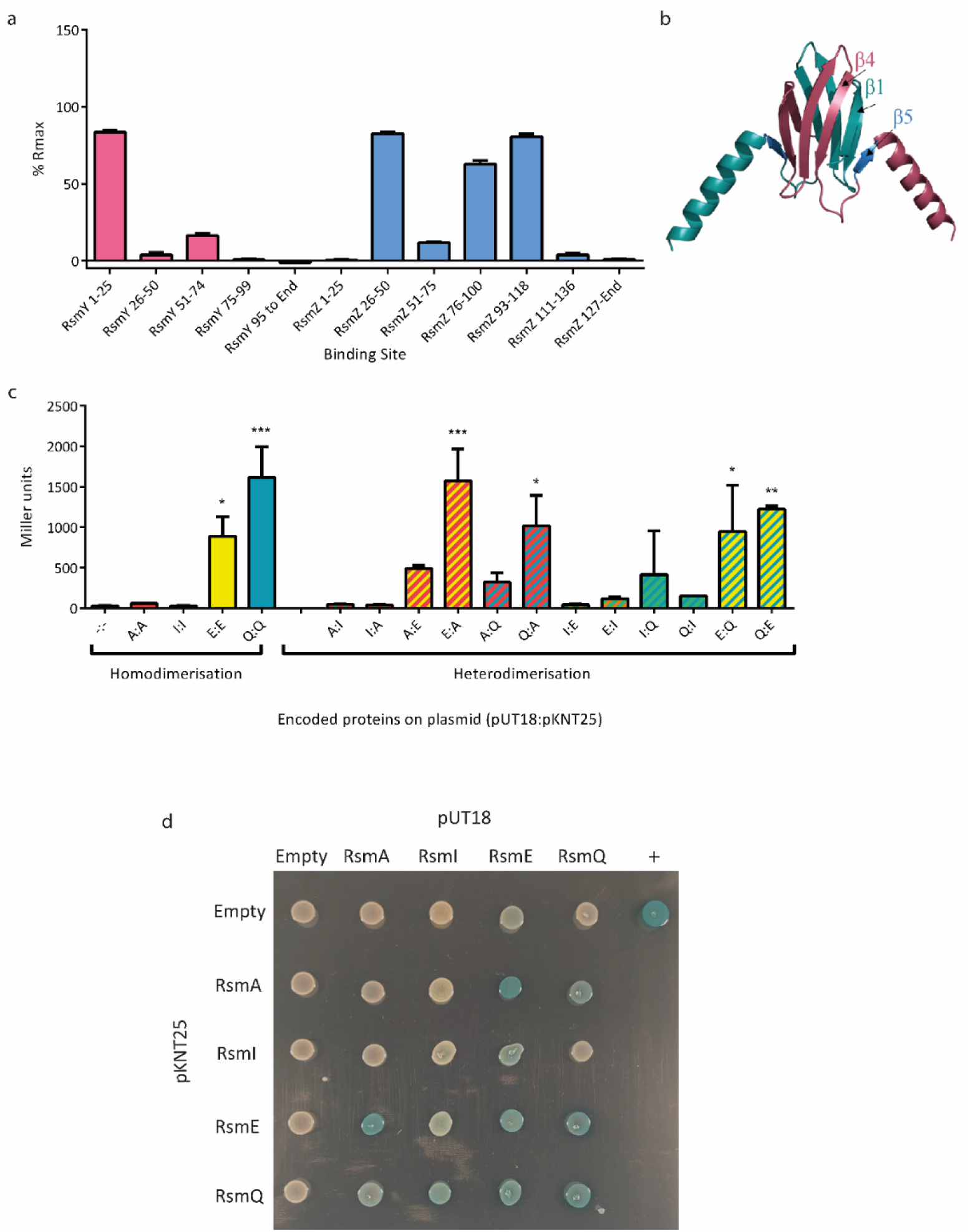
RsmQ can both homo- and heterodimerise. a) Percentage R_max_ values for RsmQ binding to portions of the ncRNAs RsmY (pink) and RsmZ (blue) showing preferential binding to ssDNAs that contained the binding site in a hairpin loop. b) AlphaFold model of RsmQ suggests that it forms homodimers (monomers shown in contrasting colours), with the RNA-binding region highlighted in marine (B5). c) Quantitative bacterial-2-hybrid β-galactosidase assays showing interactions between pUT18c and pKNT25 fusions are shown for RsmA (A), RsmE (E), RsmI (I) and RsmQ (Q). Results were analysed by a one-way ANOVA (p<0.0001) with comparisons against the Empty plasmid control (-:-) indicated (*p<0.05, ** p<0.01, *** p<0.001, ****p<0.0001). Additional controls are shown in Figure S4. d) Representative image of qualitative β-galactosidase assays on agar plates. pKT25 fusions are shown in rows and pUT18c fusions in columns, with the indicated protein/empty vector present in each case.

Rsm proteins have been seen to homodimerize and are regularly found as homodimers within the cell (21, 22, 56). With the exception of RsmN from *P. aeruginosa*, they generally have a conserved tertiary structure. Furthermore, an AlphaFold (57) model of RsmQ was shown to be highly similar to the crystal structures of SBW25 host Rsm proteins (Figure 5b)(56). We hypothesised therefore that RsmQ may also interfere with regulation by forming heterodimers with host Rsm proteins. To test this, we expressed *rsmQ* and the SBW25 host *rsm* genes heterologously in *E. coli* using the BACTH system. Interestingly, with the exception of RsmI, we saw evidence of homo- and heterodimerisation within and between the Rsm proteins. Both RsmE and RsmQ homodimerised, and heterodimerisation was observed between all pairwise combinations of RsmA, RsmE and RsmQ (Figure 5c and d). These results therefore support two indirect mechanisms for RsmQ regulation of the SBW25 proteome in addition to direct mRNA binding: either by sequestering ncRNAs, or directly interfering with the activity of host Rsm regulators.

### RsmQ causes phenotypic changes in SBW25

Given the largescale changes that RsmQ caused to the SBW25 proteome, we hypothesised that these altered protein abundances would in turn affect bacterial behaviour. To test this, we quantified the impact of RsmQ on ecologically important traits normally controlled by the Gac-Rsm regulatory system. Specifically, we initially quantified swarming motility and production of exopolysaccharide/adhesin (measured using an indirect Congo red binding assay (58)) by SBW25 in the presence and absence of *rsmQ*. To examine the direct impact of *rsmQ* on chromosomally encoded genes, *rsmQ* was expressed under an inducible promoter on a multicopy plasmid, in the absence of pQBR103^Km^. Overexpression of *rsmQ* led to a complete loss of swarming motility and a significant increase in Congo red binding (Figure 6a and c). This suggests that *rsmQ* shifts SBW25 towards a more sessile lifestyle as characterised by reduced flagellar motility and increased production of attachment factors and/or extracellular polysaccharides associated with biofilm formation.

**Figure 6:**
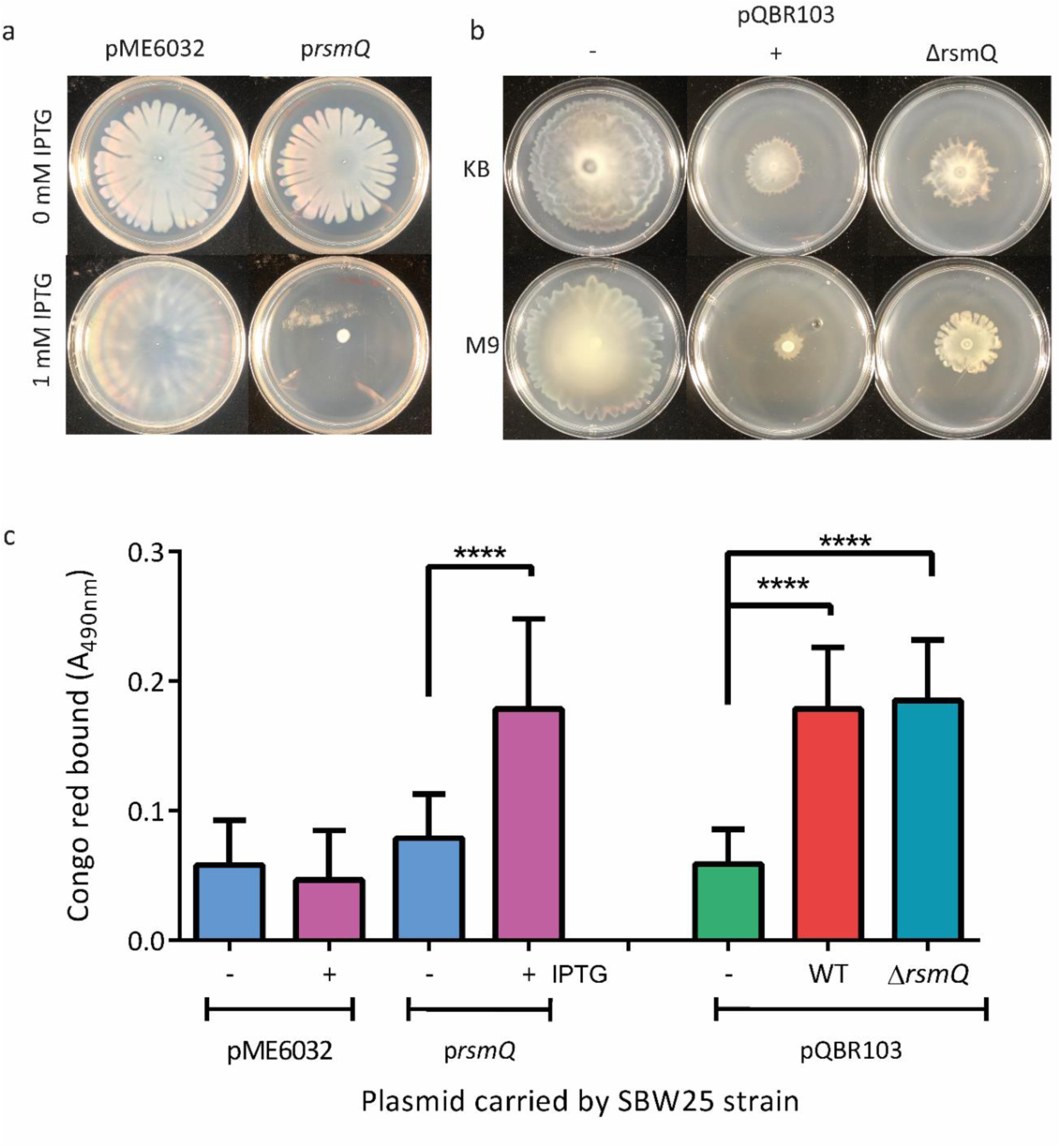
Motility and biofilm formation are impacted by RsmQ. A) 48h swarming motility assays for SBW25 containing pME6032 +/- *rsmQ.* B) 72h swarming motility assays for SBW25 cells either plasmid free (-), or carrying pQBR103^Km^ (+) or pQBR103^Km^-Δ*rsmQ* grown on 0.5% agar plates with media as indicated. C) Congo red absorbance (A_490_) of SBW25 strains after 48 hours (light blue, pink bars) or 72 hours (green, red, dark blue bars). ANOVA results show statistically significant differences for both overexpression (p< 0.0001) and deletion (p< 0.0001). Statistical significance from multiple comparisons is indicated (p<0.0001, ****).

To test if *rsmQ* had similar effects on SBW25 behaviour when encoded on pQBR103, we repeated the experiments using SBW25 with or without pQBR103^Km^ ± *rsmQ*. Acquisition of pQBR103^Km^ caused reduced swarming motility and Congo red binding relative to plasmid-free SBW25. However, deletion of *rsmQ* only partially ameliorated the reduction in swarming motility (Figure 6b) and had no effect on Congo red binding (Figure 6c). The expression of Rsm proteins is normally tightly controlled by the cell, these results suggest that at high concentrations RsmQ is able to override the native cellular control to cause drastic phenotypic changes that are not observed at the native level. This is also consistent with our proteomic data (Figure 3b), which showed little or no impact of RsmQ on the abundance of structural biofilm or motility proteins such as flagella and adhesins, but a significant impact on chemotaxis pathways.

It was therefore considered that the role of RsmQ may be in the perception and uptake of specific nutrients, and therefore any phenotypic changes may be carbon source dependent. We tested the effect of the nutrient environment on swarming motility phenotypes and observed that pQBR103 carriage strongly effected swarming motility on poorer carbon sources (Figure S3), with the loss of *rsmQ* leading to a small restoration of swarming, again suggesting *rsmQ* is involved in manipulating the cellular perception of the environment.

### Carbon source sensing by RsmQ

Rsm proteins were first characterised by their involvement in carbon storage and metabolism (e.g. CsrA in *E. coli*) and the regulation of secondary metabolism (59, 60). Although small phenotypic changes were observed on solid media (Figure S3), it remained unclear the extent to which RsmQ was involved in the sensing of specific carbon sources. The proteomic and phenotypic data suggested a role for RsmQ in sensing and responding to a variety of carbon sources.

Carbon source utilisation was compared between plasmid free SBW25 cells and cells carrying pQBR103^Km^ ±*rsmQ* using BioLog PM1 and PM2A plates, with NADH production being used as a readout. Cells were grown on 190 different carbon sources to determine the metabolic changes that occur with plasmid carriage. After 24 hours several differences were observed, with the majority of changes relating to amino acid utilisation (Table S2 and Figure 7). The pQBR103^Km^- Δ*rsmQ* containing-strain showed decreased metabolism of L-alanine, L-aspartic acid and L-arginine, each of which has also been found within root exudates (61) compared to SBW25 pQBR103^Km^. Proteomic data suggests that RsmQ represses amino acid transportation and metabolism in rich media conditions. However, when these are available as the sole carbon source, they are more easily metabolised by cells that have RsmQ present, supporting the idea that RsmQ is involved in the regulation of amino acid uptake and metabolism.

**Figure 7.**
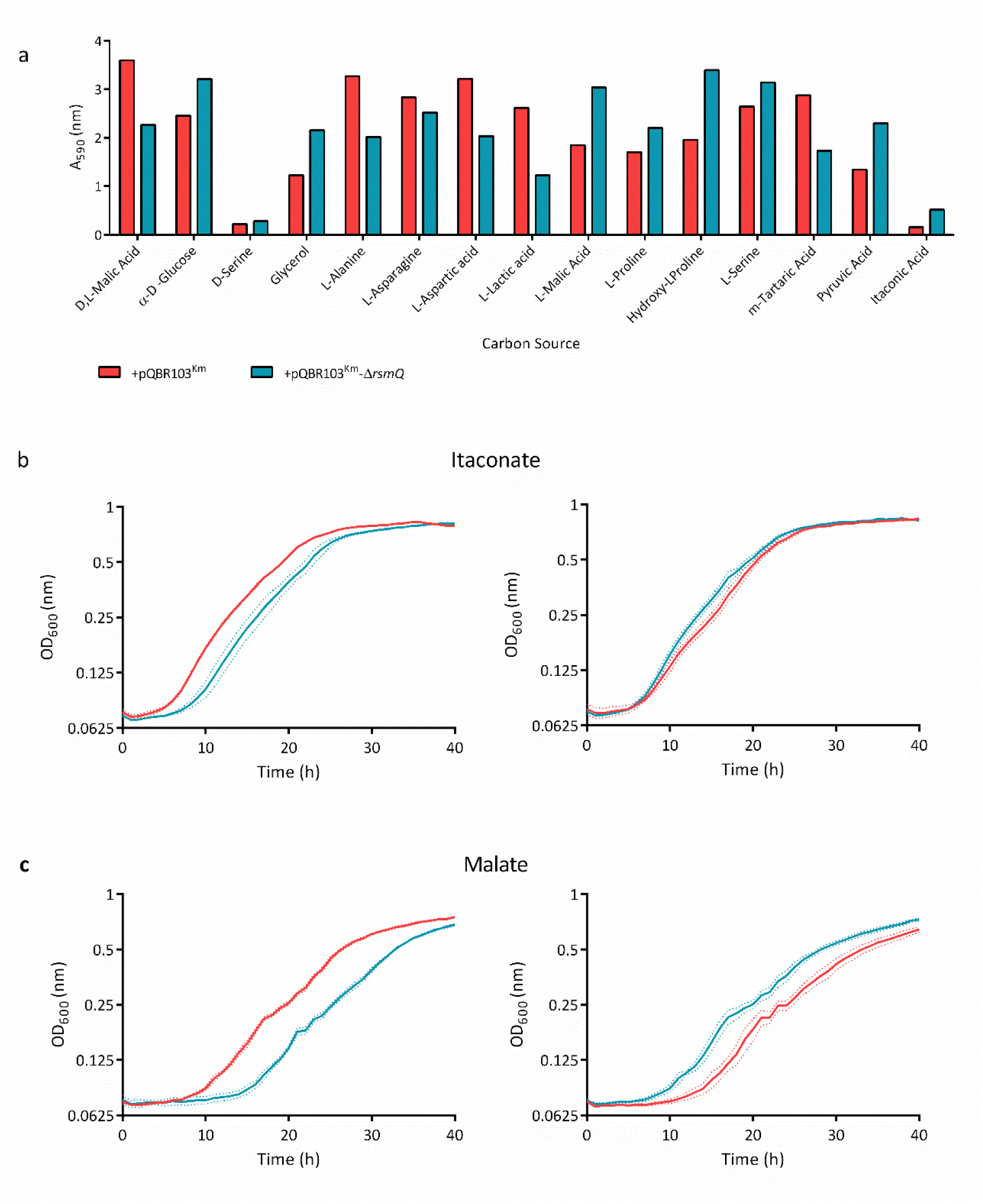
RsmQ is involved in the control of carbon source metabolism. a) Selected results from BioLog carbon source screens showing the differences in metabolism for relevant carbon sources between SBW25 cells carrying pQBR103^Km^ (red) and pQBR103^Km^-Δ*rsmQ* (blue). Representative growth curves are shown for itaconate (b) and malate (c). Cells were incubated at 28°C for 48 hours and measured at A_600_ every 30 minutes for 40 hours. Cells carrying pQBR103^Km^ are shown in red and those carrying pQBR103^Km^-Δ*rsmQ* in blue with standard deviation of three technical replicates shown as dotted lines. A minimum of six independent biological replicates were carried out for each carbon source.

As well as this, the metabolism of other carbon sources typically found in plants, such as Hydroxyl-L-Proline (62) and m-Tartaric acid (63) were impacted by the loss of RsmQ (Figure 7a). This, along with the increase in D,L-malic acid (malate) metabolism suggest that there may be a link between the carbon sources utilised by plasmid carriers and their plant host species (Figure 7a). This is particularly interesting as malate is a major component of root exudates (64, 65), suggesting that plasmid carriage may increase growth rate when in close proximity to plant roots.

To interrogate this further we selected a subset of carbon sources shown to be metabolised differently by SBW25 containing pQBR103^Km^ ±*rsmQ*. Both itaconate and malate were chosen as interesting and biologically relevant candidates that are found within root exudates (66). In M9 minimal media all tested cells were able to metabolise itaconate, in contrast to the BioLog assays, which was possibly a consequence of the different media bases used. While our M9 growth data did not simply replicate the BioLog results, itaconate and malate metabolism were both affected by *rsmQ* deletion. Curiously, RsmQ appeared to exert a biphasic effect on growth in the presence of either carbon source (Figure 7b and c). Multiple growth curves showed a difference in growth rate between pQBR103^Km^ and pQBR103^Km^-Δ*rsmQ*. However, which of these genotypes had a growth advantage (or penalty) constantly varied between different biological replicates (Figure 7b and c), suggesting that RsmQ may function to stabilise a bistable metabolic system, consistent with its chromosomal counterpart (67).

## Discussion

By encoding homologues of bacterial regulators plasmids can manipulate the expression of chromosomal genes and thereby alter the behaviour and phenotype of their host bacterial cells (4) . Previous studies of PCC have identified plasmid-encoded transcriptional regulators with limited and specific regulons (1). Here we expand the known molecular mechanisms mediating PCC to include a global post-transcriptional regulator, RsmQ. We show that RsmQ is a homologue of the widespread Csr/Rsm family of translational regulators. RsmQ has only a minimal effect on the *P. fluorescens* transcriptome but a large impact on the proteome both directly, through binding mRNAs to control their translation, and indirectly, through interactions with components of the cellular Rsm regulatory system. RsmQ alters the abundance of chemotaxis/motility and metabolism related proteins, leading to observable growth differences in distinct carbon sources.

The soil surrounding plant roots is a complex and intensely competitive environment. Rhizosphere-dwelling bacteria respond to their surroundings at an individual level using networks of signalling proteins (68) that control bacterial behaviour, enabling effective colonisation and environmental adaptation. Simultaneously, the distribution of genes in rhizosphere metagenomes are under intense selection to best fit the prevailing environment. In recent years, substantial progress has been made towards understanding both the regulatory pathways that control bacterial rhizosphere colonisation (69), and the effect of environmental inputs on microbiome composition and species’ metagenomes (70–72). Horizontal gene transfer, for example by conjugative plasmids, is well understood as an important driver of genetic adaptation ((35, 73, 74)). However, the influence of plasmid encoded regulatory genes in bacterial signalling and behaviour, and the importance of this process to bacterial fitness and evolution in the rhizosphere is much less clear.

To address these questions, we first examined the distribution of *rsm*-family genes on plasmids in the PROKKA database. Numerous plasmids were found to encode *rsm* homologs, although these were not evenly distributed. Plasmids associated with some taxa, including Pseudomonadaceae, frequently contained *rsm* genes, while those associated with others had none. This distribution suggests that plasmid encoded Rsm proteins may fulfil important functional roles that are associated with particular groups of microbes. Plasmid carriage of *rsm* regulators appears to be a convergent trait that has emerged many times, potentially in response to similar environmental pressures affecting different plasmids. Overall, our analysis suggests that diverse plasmids have acquired *rsm* regulator genes from their bacterial hosts over time, and that these genes now appear to be evolving distinctly from their chromosomally located ancestors. Interestingly, *rsm* genes have also been identified on bacteriophage (38), suggesting that diverse mobile genetic elements may exploit host post-transcriptional regulation to ultimately serve the fitness interests of mobile genetic elements.

Next, to determine the functional role of plasmid borne Rsm regulators we examined the pQBR103-encoded protein RsmQ, in the plant-associated rhizobacterium *P. fluorescens* SBW25. Our data shows that RsmQ functions as a global, post-transcriptional regulator of metabolism, nutrient transport and chemotaxis pathways. In this respect RsmQ-linked PCC differs markedly from most recently described PCC systems, which involve altered transcription of a small number of genes upon plasmid carriage, typically relating to one or two bacterial phenotypes (1). By contrast, RsmQ-driven regulation occurs predominantly at a post-transcriptional level and is global in scope, affecting the abundance of hundreds of proteins and extensively subverting bacterial motility and metabolic pathways to the benefit of the plasmid. Our results implicate RsmQ and by extension other, similar plasmid-borne regulators, in considerably more extensive control of bacterial behaviour than has previously been observed.

In contrast to earlier studies (45),we saw relatively little effect of plasmid carriage or *rsmQ* on SBW25 gene transcription. In itself, this finding is not particularly surprising as our test conditions were chosen to interrogate RsmQ function and differed markedly from those used previously. Nonetheless, this does suggest that the nature and extent of pQBR103 control of SBW25 transcription is highly dependent on the surrounding environment. Overall, this suggests that RsmQ appears to reprogram chemotaxis and metabolism towards a more sessile, biofilm forming lifestyle. This is consistent with the phenotypic data for pQBR103 carriage, where a reduction in swarming motility is partially recovered by *rsmQ* deletion, as well as changes in both bacterial growth in a carbon source dependent manner.

RsmQ functions in SBW25 by interacting with the host Gac/Rsm pathway in several different ways. Firstly, RsmQ binds to specific RNA target sequences at conserved binding motifs in a similar manner to chromosomally encoded Rsm proteins. RsmQ has been shown to have a binding preference for both short (GGA) and extended (AnGGA) motifs, especially where these are presented on the loops of ssDNA hairpins. To some extent, this RNA binding effect contributes to RsmQ-mediated PCC by directly regulating gene targets through chromosomal mRNA binding. In support of this, we confirmed direct RsmQ binding to ssDNA sequences corresponding to the upstream regions of several RsmQ-regulated genes. Furthermore, the AnGGA RsmQ binding motif is only found upstream of around half of RsmQ-affected SBW25 genes. Thus, direct mRNA binding probably only accounts for a fraction of the observed RsmQ regulon.

At least part of the observed RsmQ regulon is likely the result of indirect proteomic adaptation in response to changes induced by RsmQ. This has been shown previously for the translational regulator Hfq in SBW25, where the direct targets of translational control do not correspond neatly to the observed proteomic regulon (5, 55). Another, potentially direct route for RsmQ function is through interaction with the host ncRNAs RsmZ and RsmY. RsmQ binds strongly to several RsmY/Z motifs as shown by SPR, suggesting that similar binding takes place in the SBW25 cell. This binding interaction could in-turn mediate pleiotropic changes in gene translation, either through RsmY/Z titration of RsmQ away from its mRNA binding targets, or alternatively via a reduction in the amount of RsmY/Z available to modulate the activity of the host Rsm proteins. Both of these alternative, and potentially antagonistic mechanisms may function simultaneously to some extent, with the abundance of RsmQ and RsmY/Z in the cell determining their relative importance to cellular Rsm function.

In addition to its interaction with host RNAs, we also saw evidence for extensive Rsm protein homodimerisation, as well as heterodimerisation between RsmQ and the host proteins RsmA and RsmE. Rsm protein homodimerisation is a well-characterised trait, with structural evidence supporting widespread homodimer formation in *Pseudomonas* spp. (21, 56, 75). However, to our knowledge Rsm proteins have not previously been shown to form heterodimers. The mechanistic consequences of RsmQ/A/I heterodimerisation are currently unknown. The simplest explanation is that the three Rsm proteins are functionally and mechanistically interchangeable, and Rsm heterodimers have no *in vivo* function, however this seems unlikely given their distinct regulons (18). Alternatively, RsmQ may have an agonistic or antagonistic effect on the host Rsm proteins, increasing or reducing their RNA binding affinity. Heterodimerisation may also influence mRNA binding preference, shifting the global Rsm regulatory response towards an outcome that benefits horizontal or vertical plasmid transmission, although this is yet to be determined.

The presence of multiple, interconnected layers of regulation make signalling phenotypes challenging to interrogate in a laboratory setting. For example, Hfq is global master regulator of translation (7, 55) and has an overlapping regulon with the Rsm proteins (76) as well as circumstances in which Hfq and RsmA directly interact (77), adding further regulatory complexity. Hfq controls multiple phenotypes in SBW25 (7), however the effects of Hfq dysregulation upon *rsmQ* deletion appeared to be modest, particularly when compared to secondary Hfq control by the RimABK system, for example (7, 78). An intriguing possibility is that the plasmid-borne Hfq homologue pQBR0137, whose abundance increases in the absence of *rsmQ*, may compensate for reduced chromosomal Hfq activity. The reasons behind any potential compensatory regulation are unknown and the subject of active investigation, although it is striking that multiple plasmid encoded regulators may in fact be functioning in a coordinated manner in SBW25, modulating cellular responses to the environment at a global level.

Carriage of pQBR103^Km^ leads to an increase in biofilm formation and reduced motility, two Gac-system associated traits. From the perspective of the plasmid this makes sense; tightly packed biofilms are more likely to support plasmid transmission (4). These key phenotypes associated with plasmid carriage were also seen upon *rsmQ* overexpression. However, while *rsmQ* deletion from pQBR103 partially ameliorated the plasmid-induced loss of motility, we saw no effect on biofilm, suggesting that this *rsmQ* overexpression phenotype may be non-specific. This could potentially be explained by excess RsmQ binding to the *fleQ* mRNA, as FleQ is a master regulator of *Pseudomonas* behaviour that controls both motility and biofilm formation (16, 79). Gac/Rsm is a tightly controlled global regulatory system, with multiple links to other signalling pathways and extensive in-built functional redundancy. Therefore, the relatively subtle phenotypic differences seen upon *rsmQ* deletion under lab conditions are unsurprising.

The effects of RsmQ on SBW25 metabolism and growth were strikingly inconsistent. Across repeated, independent assays *rsmQ* conferred either a growth advantage or a penalty, apparently at random. This suggests that the outcome of assay was defined stochastically, and implies a degree of bistability within the system. Regulatory bistability has been shown for several bacterial systems (80, 81) and has been linked to Gac/Rsm (67, 76). It is possible therefore that the loss of regulatory control by RsmQ has knock-on effects within the wider regulatory network, inducing stochastic, bistable growth phenotypes compared to WT pQBR103^Km^.

In conclusion, we propose that RsmQ functions as a global PCC regulator that both directly controls host mRNA translation and interferes with the host Gac/Rsm pathway and the master regulator Hfq. RsmQ activity remodels host metabolism and suppresses motility as part of a wider PCC programme that directs *P. fluorescens* towards biofilm formation and a sessile lifestyle, where horizontal plasmid transmission is likely to be favoured. We hypothesise that core elements of this regulatory paradigm will be shared between diverse plasmid-borne Rsm proteins, with the PCC regulon tuned in each case to best support plasmid fitness in the host environment. More broadly, whereas plasmid accessory genes are often conceptualised as directly providing distinct, novel functions such as toxin efflux or enzymatic degradation of metabolic substrates, out work shows that plasmids might also find success, and indeed exert ecologically important effects, by manipulating and tuning the expression of functions encoded by genes already resident within the cell.

## Materials and Methods

### Strains and growth conditions

Strains and plasmids are listed in Table S3 and primers are listed in Table S4. Unless otherwise stated *P. fluorescens* SBW25 were grown at 28°C and *E. coli* strains at 37°C in Lysogeny broth (LB) (82) solidified with 1.5% agar where appropriate. Liquid cultures were grown in 10 mL microcosms at 28°C for *P. fluorescens* and 37°C for *E. coli* at 250 rpm unless otherwise stated. Minimal media was made using M9 salts supplemented with 2 mM MgSO_4_ and 0.1 mM CaCl_2_ and each carbon source present at 0.4%. For motility assays plates were solidified with 0.5% agar. Gentamicin (Gent) was used at 25 µg ml^-1^, Streptomycin (Strep) at 250 µg ml^-1^, Kanamycin (Kan) at 50 µg ml^-1^, Carbenicillin (Carb) at 100 µg ml^-1^, Tetracycline (Tet) at 12.5 µg ml^-1^, IPTG at 1 mM and X-gal at 40 µg ml^-1^.

### Molecular biology procedures

Cloning was carried out in accordance with standard molecular biology techniques. All pTS1 plasmid inserts were synthesised and cloned into pTS1 by Twist Bioscience. The ORF of *rsmQ* was amplified by PCR with primers RsmQ_EcoRI_F and RsmQ_XhoI_R and ligated between the EcoRI and XhoI sites of pME6032. The ORF of *rsmQ* with a TEV cleavage site and a hexahistidine tag at the C-terminus was synthesised by Twist Bioscience. The ORFs of each Rsm protein were amplified by PCR using the primers indicated in Table S3. The fragment in each case was cloned between the NdeI and XhoI sites of pET28a. Bacterial-2-hybrid plasmids were made by Gibson assembly (RsmE/I) and restriction cloning into the BamHI and EcoRI sites of pKNT25 and pUT18C.

### Transformation of *Pseudomonas* strains

Overnight cultures of each strain were grown in LB media at 28°C, 250 rpm shaking then harvested at 6000 x*g* for 8 minutes. Cell pellets were washed twice with 0.3 M sucrose, then the pellet was resuspended in a final volume of 150 µL and placed in a 2 mm electroporation cuvette with either 2 µL of replicative plasmids or 5 µL of integrative plasmids (60-100 ng/µL concentration) and incubated at RT for 2 minutes. Cells were electroporated at 2.5 kV and recovered in 3 mL LB medium at for 3 hours before being plated onto LB agar containing the appropriate antibiotic. Plates were incubated for 24-48 hours at 28°C and transformed colonies re-streaked onto fresh selective media.

### Conjugations of pQBR103^Km^

Donor (*P. fluorescens* SBW25 ΩStrepR-LacZ + pQBR103^Km^ and *P. fluorescens* SBW25 ΩStrepR-LacZ + pQBR103^Km^ Δ*rsmQ*) and recipient strains (*P. fluorescens* SBW25 ΩGentR WT) were plated onto their respective selective antibiotics. Overnight cultures were set up in LB medium for each of the strains and grown overnight. 10 mL glass microcosms of Kings Broth (KB) medium were inoculated with 20 µL of the donor strain and 80 µL of the recipient strain and incubated overnight without shaking. 50 µL of this overnight culture was plated onto LB medium supplemented with Gent and Kan.

### Allelic Exchange

Deletion constructs were created by Twist bioscience and extracted from *E. coli* DH5-α cells. *P. fluorescens c*ells were transformed as above and incubated for 48 hours until colonies appeared. Colonies were re-streaked to single colonies on fresh LB-Tet agar and incubated for 24 hours. A single colony was picked and grown overnight in 50 mL of LB medium (containing Kanamycin for pQBR103^Km^ allelic exchange). The culture was serially diluted, and the 10^-5^ to 10^-8^ dilutions were plated onto LB agar plates containing 10% sucrose (and kanamycin for pQBR103^Km^ allelic exchange). Single colonies on sucrose plates were checked for tetracycline sensitivity and confirmed as mutants by PCR.

### Swarming motility assays

Motility plates were made with 20 mL of sterile 0.5% agar in M9 GC (M9 minimal media supplemented with 0.4% Glucose and 0.4% Casein amino acids) media unless otherwise indicated and dried for 1 hour in a laminar flow hood, rotated 180 degrees after 30 minutes. 3 µL of an overnight culture adjusted to an OD_600_ = 1 was spotted onto the centre of the plate and the lid replaced. Plates were incubated face up at room temperature for 72 hours undisturbed and then imaged. For overexpression strains, filter sterilised 0.5 mM IPTG and tetracycline were added to the induced samples.

### Congo red binding assays

Cultures of each strain were grown overnight in LB microcosms with selection. 10 µl spots were placed onto 20 mL KB agar plates and incubated for 72 hours. For overexpression strains, filter sterilised 0.5 mM IPTG and tetracycline were added to the induced samples. Colonies were removed from the plate and placed into 1 mL of 0.003% sterile Congo red solution and incubated at 37°C, 200 rpm shaking for 2 hours. Cell material was removed by centrifugation (13,000 x*g* for 5 minutes) and absorbance was measured at 490 nm using a SPECTROstar nano plate reader (BMG).

### BioLog carbon source screening

Two colonies were picked from freshly streaked LB plates and resuspended into IF-0 inoculating fluid as per the manufacturer’s instructions. PM1 and PM2A plates were inoculated with 100 µl of inoculum and incubated at 28°C for 24 hours. Plates were imaged and the absorbance at 590 nm was read on an EON microplate reader.

### Growth rate assays

Cultures of each strain were grown overnight in LB microcosms with selection. Cells were harvested at 8,000 x*g* and washed in M9 media without a carbon source twice. Cells were then resuspended in M9 media with each carbon source (0.4% w/v) at a starting OD_600_ of 0.01 in a 96-well plate. Measurements were taken every 30 minutes for 40 hours on a FLUOstar nano plate reader (BMG) with the plate being incubated at 28°C and shaken for 2 seconds before each reading.

### Bacterial 2 hybrid assays

The ORF of RsmA/E/I/Q were cloned into pKNT25 and pUT18C using either conventional restriction enzyme cloning or Gibson assembly using standard manufacturers protocols as indicated in Table S4. Chemically competent BTH101 cells were co-transformed with both a pUT18 and a pKNT25 plasmid containing the ORF of the protein of interest using the heat shock method. Briefly, cells were incubated on ice with the plasmids for 30 minutes, followed by a 45s incubation at 42°C followed by 5 minutes on ice. Cells were recovered in 6 volumes of SOC media for one hour and plated onto LB agar supplemented with Carb, Kan and 0.5 mM IPTG. 5 mL LB+Carb+kan microcosms were grown overnight at 28°C. 100 μL of this overnight culture was used for the beta-gal assay and 5 μL spots were placed onto LB+Carb+Kan+X-gal+IPTG plates and incubated overnight.

### β-galactosidase assays

*E. coli* BTH101, 5 mL microcosms of cells carrying both pUT18C and pKNT25 plasmids were grown at 28°C. 100 μL of this was taken and incubated with 900 μL lysis buffer (60 mM Na_2_HPO_4_.7H_2_O, 40 mM NaH_2_PO_4_.H2O, 10 mM KCl, 1 mM MgSO_4_, 7.7 mM β-mercaptoethanol, 0.001% SDS) and 20 μL chloroform at 28°C for >10 minutes until cells are lysed. 200 μL of 4 mg mL^-1^ ONPG was added, and samples monitored until the substrate had turned yellow. To stop the reaction 500 μL of 1M Na_2_CO_3_ was added, and the absorbance was taken at 420 and 550 nm using a FLUOstar plate reader (BMG) and OD_600_ of each sample was measured using a spectrophotometer. The Miller units were calculated using the standard calculation (82).

### Protein purification

The ORF of *rsmQ* was synthesised with a C-terminal TEV cleavage site extension and a 6x Histidine tag (Twist bioscience) and cloned into pET29a between the NdeI and XhoI sites. The plasmid was transformed into BL21(DE3) pLysS (Promega) by heat shock. 2.5 L of culture was inoculated at 1:50 from an overnight culture and grown until mid-log phase (OD_600_ = ∼0.6). Cultures were induced with 1 mM IPTG and grown for 16 hours at 37°C. Cells were harvested at 6000 x*g*, 4°C for 15 minutes and resuspended in binding buffer (20 mM Tris-HCl, 500 mM NaCl, 10 mM imidazole, 5% glycerol, pH 7.5) containing 1 mg ml^-1^ lysozyme (sigma), 1 complete protease inhibitor tablet EDTA-free and 5 µl DNaseI (Promega), lysed using a cell disruptor and the insoluble fraction removed by centrifugation (15,000 x*g*, 25 minutes, 4°C). The soluble fraction was loaded onto a HisTrap HP 5 mL column (Cyvitia) and washed with binding buffer (20 mM Tris-HCl, 500 mM NaCl, 2.5% glycerol, pH 7.5) with 50 mM Imidazole to remove non-specific contaminants. Proteins were eluted over a gradient of 50-500 mM Imidazole and fractions analysed by SDS-PAGE.

Fractions containing RsmQ were dialysed overnight at 4°C into SEC buffer (50 mM Tris-HCl, 200 mM NaCl, 2.5% Glycerol, pH 7.5) and further purified by size exclusion chromatography using an Superdex S75 column (Cyvitia). Fractions were analysed by SDS-PAGE and pure fractions were concentrated to 3 mg ml^-1^ and stored at -80°C until needed.

For H43A and R44A plasmids were created by site-directed mutagenesis PCR using overlapping primers (RsmQ_H43A_F, RsmQ_H43A_R, RsmQ_R44A_F, RsmQ_R44A_R), confirmed by sequencing and purified as for the WT.

### Surface plasmon resonance

Single stranded DNA oligos (ssDNA) with the ReDCaT linker region at the 3’ end were synthesised by IDT and diluted to a final concentration of 1 mM in HBSEP+ buffer (10 mM HEPES, 150 mM NaCl, 3 mM EDTA and 0.05% v/v Tween 20, pH 7.4). All primer sequences can be found in Table S4 (ReDCaT). RsmQ, RsmQ H43A and RsmQ R44A were diluted to 1000 and 100 nM concentrations in HBSEP+ buffer. ssDNA oligos were synthesised (IDT) and diluted to a final concentration of 1µM in HBSEP+. SPR measurements were recorded at 20 °C using a Biacore 8k system using a ReDCaT SA sensor chip (GE Healthcare) with 8 immobilised channels as described in (43). RsmQ interaction was first analysed using an affinity method to examine presence and absence of binding to each of the oligos.

### RNAseq

5 independent conjugations were set up for each biological replicate as described above. A single colony from each conjugation event was picked and grown overnight in 10 mL KB medium with Gent and Kan at 28°C, 230 rpm. The OD_600_ of these cultures was measured and 60 mL KB cultures were set up at OD_600_ = 0.01. Cultures were grown at 28°C, 230 rpm until OD_600_ = 1.4. 2 mL of this culture was harvested for RNA extraction and 50 mL was taken for whole proteome analysis. Pellets were collected at 8000 x*g*, 10 minutes at 4°C and flash frozen in liquid nitrogen before storage at -80°C.

For RNAseq the pellets were resuspended in 150 µL 10 mM Tris-HCl pH 8 and mixed with 700 µL of ice cold RLT+BME (RLT buffer (Qiagen) supplemented with 1% β-mercaptoethanol) and cells were lysed using a Fastprep (MP Bio) using Lysis matrix B beads (MP Bio). Lysis matrix was removed by centrifugation (13,000 x*g*, 3 minutes) and the supernatant was added to 450 µL of ethanol. The supernatant was applied to a RNeasy column and RNA extraction was performed as per the manufacturer’s instruction including the on-column DNA digest. After extraction a Turbo DNase (Promega) digest was performed as per the manufacturer’s instruction and total RNA yield was quantified using a Qubit RNA broad spectrum assay kit as per the manufacturer’s instructions. Library construction, rRNA depletion and paired-end Illumina sequencing (Novaseq 6000, 2x150 bp configuration) was performed by Novogene. Reads provided by Novagene (as fastq.gz files) were mapped to the genome of Pseudomonas fluorescens (NCBI accession AM181176.4) and the plasmid pQBR103 (NCBI accession AM235768.1), using the “subread-align” command of the Subread package (83). The resulting .bam files were then sorted and indexed using the appropriate functions from the Samtools package (84). A custom Perl script was used to make a saf file for all the gene in the genome and the plasmid. The “featureCounts” tool of the Subread package was used to count the reads mapping to every gene. The counts were read into a DGEList object of the Bioconductor package edgeR and a quasi-likelihood negative binomial generalized log-linear model was fitted to the data using the “glmQLFit” function of edgeR(85). Genewise statistical tests were conducted using the “glmQLFTest” function of edgeR. Finally, the “topTags”. Processed data is deposited in ArrayExpress (E-MTAB-11868).

### Whole proteome analysis

Cells were grown as above and stored at -80°C until the proteome was extracted. Three samples were thawed on ice and resuspended in ice cold 500 µL Lysis buffer (20 mM Tris, 0.1 M NaCl, pH 8 + 1 complete protease inhibitor tablet). Cells were lysed by sonication at 12 mA (1 second on, 3 seconds off for 20 cycles). Insoluble material was removed by centrifugation at 4°C, 4000 x*g*, 10 minutes. The supernatant was taken and the proteome precipitated for 10 minutes with the addition of 8 volumes of acetone at RT. The proteome was pelleted by centrifugation at 7000 x*g* for 10 minutes and washed once more with acetone. Protein pellets were resuspended in 400 µl of 2.5% sodium deoxycholate (SDC; Merck) in 0.2 M EPPS-buffer (Merck), pH 8.5, and vortexed under heating for a total of three cycles. Protein concentration was estimated using a BCA assay and approx. 200 µg of protein per sample was reduced, alkylated, and digested with trypsin in the SDC buffer according to standard procedures. After the digest, the SDC was precipitated by adjusting to 0.2% TFA, and the clear supernatant subjected to C18 SPE (OMIX tips; Agilent). Peptide concentration was further estimated by running an aliquot of the digests on LCMS (see below). TMT labelling was performed using a TMTpro™ 16plex kit (Lot WB314804, Thermofisher Scientific) according to the manufacturer’s instructions with slight modifications; approx. 100 µg of the dried peptides were dissolved in 90 µl of 0.2 M EPPS buffer (MERCK)/10% acetonitrile, and 250 µg TMT reagent dissolved in 22 µl of acetonitrile was added. Samples were assigned to the TMT channels.

After labelling, aliquots of 1.5 µl from each sample were combined in 500 µl 0.2% TFA, desalted, and analysed on the mass spectrometer (see below) to check labelling efficiency and estimate total sample abundances. The main sample aliquots were then combined correspondingly to roughly level abundances and desalted using a C18 Sep-Pak cartridge (200 mg, Waters). The eluted peptides were dissolved in 300 µl 0.1% TFA and fractionated with the Pierce™ High pH Reversed-Phase Peptide Fractionation Kit (Thermo) according to the manufacturer. Fractions for the mass spectrometry analysis were eluted sequentially with the following concentrations of acetonitrile: 7,5%, 10%, 12.5%, 15%, 17.5%, 20%, 30%, 40%, 50% and dried down and resuspended in 0.1 % TFA, 3% acetonitrile.

Aliquots were analysed by nanoLC-MS/MS on an Orbitrap Eclipse™ Tribrid™ mass spectrometer coupled to an UltiMate® 3000 RSLCnano LC system (Thermo Fisher Scientific, Hemel Hempstead, UK). The samples were loaded onto a trap cartridge (PepMap 100, C18, 5um, 0.3x5mm, Thermo) with 0.1% TFA at 15 µl min^-1^ for 3 min. The trap column was then switched in-line with the analytical column (nanoEase M/Z column, HSS C18 T3, 1.8 µm, 100 Å, 250 mm x 0.75 µm, Waters) for separation using the following gradient of solvents A (water, 0.1% formic acid) and B (80% acetonitrile, 0.1% formic acid) at a flow rate of 0.2 µl min^-1^ : 0-3 min 3% B (parallel to trapping); 3-10 min linear increase B to 8 %; 10-90 min increase B to 37%; 90-105 min linear increase B to 50 %; followed by a ramp to 99% B and re-equilibration to 3% B. Data were acquired with the following parameters in positive ion mode: MS1/OT: resolution 120K, profile mode, mass range m/z 400-1800, AGC target 100%, max inject time 50 ms; MS2/IT: data dependent analysis with the following parameters: top10 in IT Rapid mode, centroid mode, quadrupole isolation window 0.7 Da, charge states 2-5, threshold 1.9e4, CE = 30, AGC target 1e4, max. inject time 50 ms, dynamic exclusion 1 count for 15 sec mass tolerance of 7 ppm; MS3 synchronous precursor selection (SPS): 10 SPS precursors, isolation window 0.7 Da, HCD fragmentation with CE=50, Orbitrap Turbo TMT and TMTpro resolution 30k, AGC target 1e5, max inject time 105 ms, Real Time Search (RTS): protein database *Pseudomonas fluorescens* SBW25 (uniprot.org, 02/2016, 6388 entries), enzyme trypsin, 1 missed cleavage, oxidation (M) as variable, carbamidomethyl (C) and TMTpro as fixed modifications, Xcorr = 1, dCn = 0.05.

The acquired raw data were processed and quantified in Proteome Discoverer 2.4.1.15 (Thermo) using the incorporated search engine Sequest HT and the Mascot search engine (Matrix Science, London, UK; Mascot version 2.8.0). The processing workflow included recalibration of MS1 spectra (RC), reporter ion quantification by most confident centroid (20 ppm), fasta databases P. fluorescens SBW25 (as for RTS) and common contaminants, precursor/fragment tolerance 6 ppm/0.6 Da, enzyme trypsin with 1 missed cleavage, variable modification was oxidation (M), fixed were carbamidomethyl (C) and TMTpro 16plex. The consensus workflow included the following parameters: unique peptides (protein groups), intensity-based abundance, TMT channel correction values applied (WB314804), co-isolation/SPS matches thresholds 50%/70%, normalisation on total peptide abundances, protein abundance-based ratio calculation, missing values imputation by low abundance resampling, two or three replicates per sample (non-nested), hypothesis testing by t-test (background based), adjusted p-value calculation by BH-method. Experimental data is deposited in ProteomeXChange (PXD033640).

### Bioinformatics and sequence analysis

The COMPASS database (37) was downloaded from https://github.com/itsmeludo/COMPASS and annotated using PROKKA 1.14.6 (86) using the default settings. PROKKA uses BLAST+ matches with the curated UniProtKB (SwissProt) databases to annotate proteins, followed by hidden Markov model (HMM)-based searches. To compare chromosomal and plasmid *csrA/rsmA* genes, chromosomal sequences associated with the COMPASS plasmids were identified and downloaded using NCBI **elink** and **efetch** tools and were reannotated using PROKKA to ensure comparability between plasmid and chromosomal sequences. *csrA/rsmA* sequences which were identical at the nucleic acid level were removed before conducting analyses (there were no identical sequences between chromosomes and plasmids). Start codons were unified across CsrA/RsmA homologues by manual examination and sequence editing. Sequences were aligned by codon alignment in PRANK v.170427 (87) using the default settings. Initial alignments had a high proportion (>70%) of gaps owing to sequence divergence towards the 3’ end of the gene, which interfered with phylogenetic analysis. Trees were therefore built using information from conserved sites, by removing columns from the alignment that consisted of majority (>60%) gaps. Duplicate sequences were removed. Trees were estimated using RAxML 8.2.12 (88) with the settings **-f a - m GTRCAT -p 12345 -x 12345 -# 100**. Qualitatively similar outcomes were obtained if gap columns and/or duplicate sequences were retained. Jukes-Cantor distance matrices were extracted from the alignments for analysis using the EMBOSS 6.6.0.0 and distances were compared using Bonferroni-corrected Wilcoxon tests. The script CSRA_TARGET.pl (89) was adapted to predict binding sites for *csrA/rsmA* in 5’ untranslated regions of the PROKKA-predicted plasmid genes, and differences in distributions of number of sites/CDS were compared between *csrA/rsmA*-encoding and non-encoding plasmids using two-sample Kolmogorov-Smirnov tests. Analyses were performed in R (4.1.2, R Core Team, Vienna, Austria), bash, and Python 3.6 within the RStudio IDE (RStudio Team, Boston, USA) with the assistance of tidyverse (90) and ggtree (91) packages. Example scripts and analyses can be found at www.github.com/jpjh/PLASMAN_RsmQ

## Acknowledgments

The authors would like to thank Clare Stevenson and Julia Mundy for their help and advice with initial SPR experiments and analysis. JGM and CMAT were supported by BBSRC Responsive mode Grant BB/R018154/1 to JGM. JGM and RHL were supported by BBSRC Institute Strategic Programme Grant BBS/E/J/000PR9797 to the John Innes Centre. AP was supported by UKRI-BBSRC Grant BB/T004363/1 to the John Innes Centre. JPH was supported by BB/R014884/1. This research was funded by the Biotechnology and Biological Sciences Research MAB, SF and SMB were supported by BBSRC grants BB/R014884/1, BB/R014884/2, BB/R018154/1 and NERC grants NE/R008825/1, NE/R008825/2. EH is supported by a NERC Independent Research Fellowship NE/P017584/1.

---

**Figure S1:**
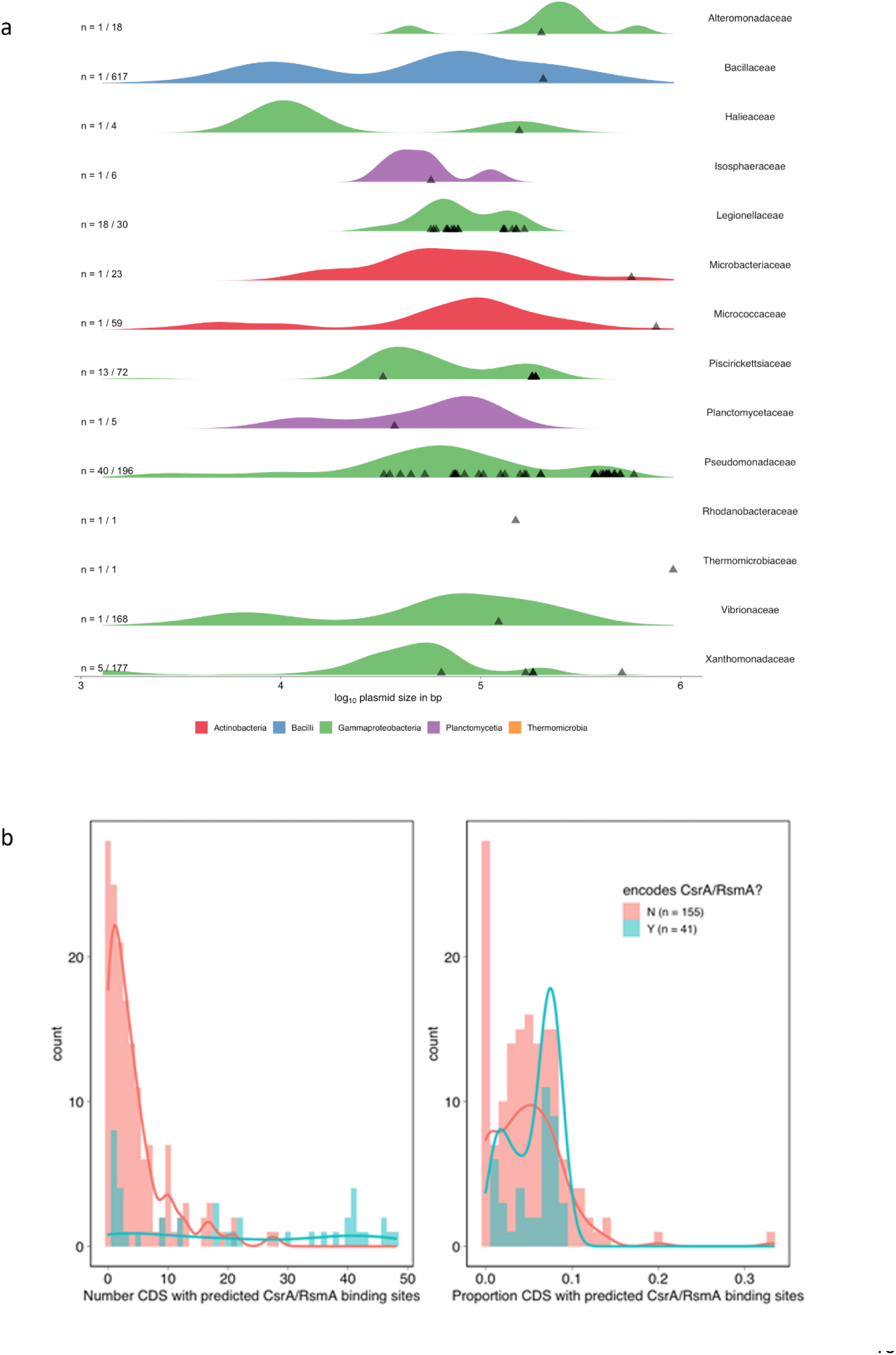
a) Across Families, CsrA/RsmA-encoding plasmids are relatively large. Size density plots for all Families with >20 plasmids and ≥1 plasmid-encoded CsrA/RsmA homologue. Each row describes a different Family. Semi-transparent triangles indicate the size of CsrA/RsmA-encoding plasmids. On the left, the proportion of total plasmids encoding CsrA/RsmA homologues for that Family. b) Comparison of putative CsrA/RsmA-regulated gene frequencies between CsrA/RsmA-encoding and non-encoding Pseudomonadaceae plasmids. Plots show overlayed histograms and density plots. Left hand plot shows absolute numbers of putative CsrA/RsmA-regulated genes, whereas right hand plot shows as a proportion of total CDS on that plasmid. Distributions were significantly different between plasmid types in both panels (Kolmogorov-Smirnov test, p < 0.001 for absolute counts, p = 0.012 for proportions).

**Figure S2:**
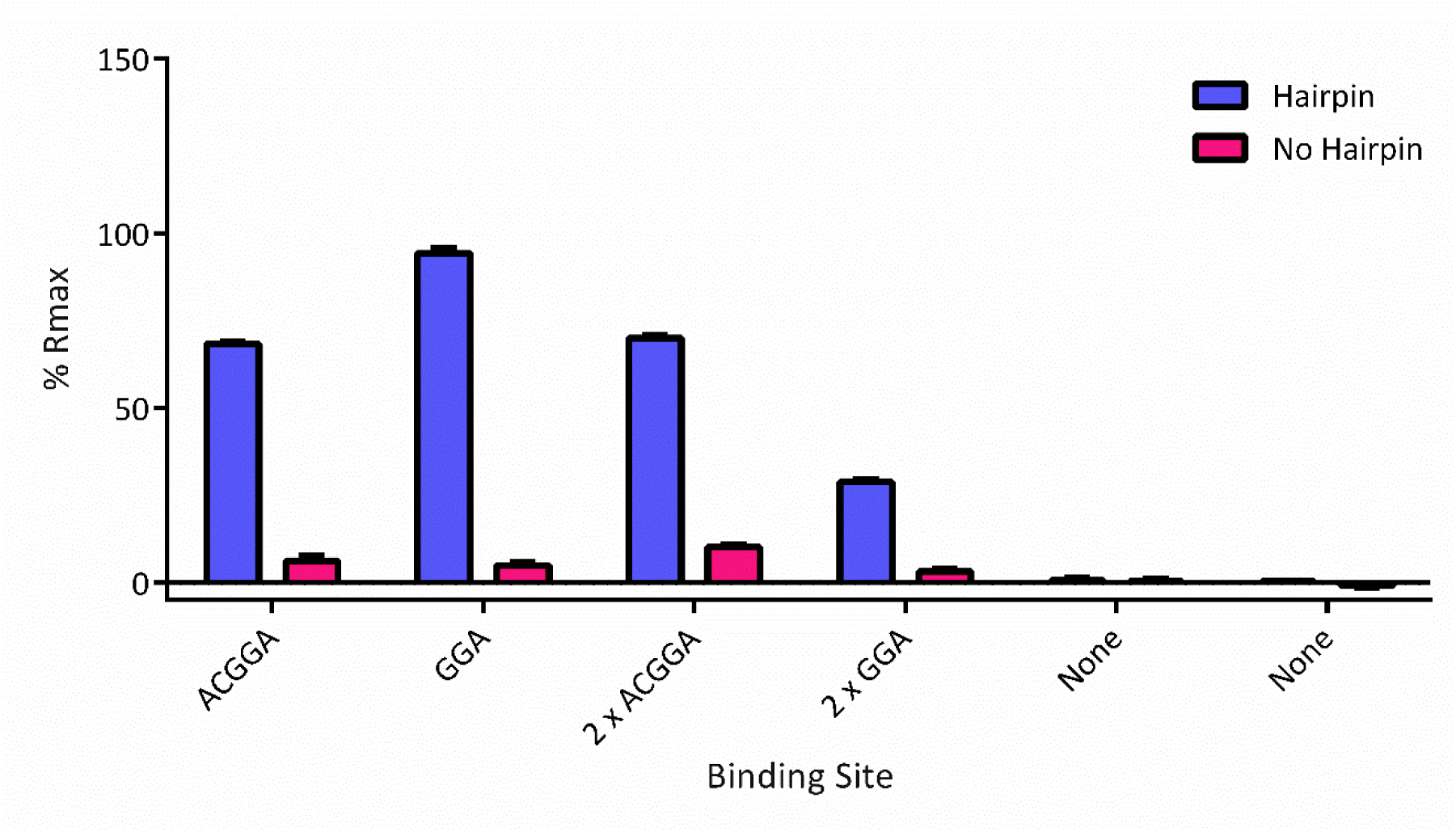
Percentage R_max_ values for RsmQ binding to ssDNAs containing the indicated binding site sequence in either a linear format (pink) or at the top of a hairpin loop (purple).

**Figure S3:**
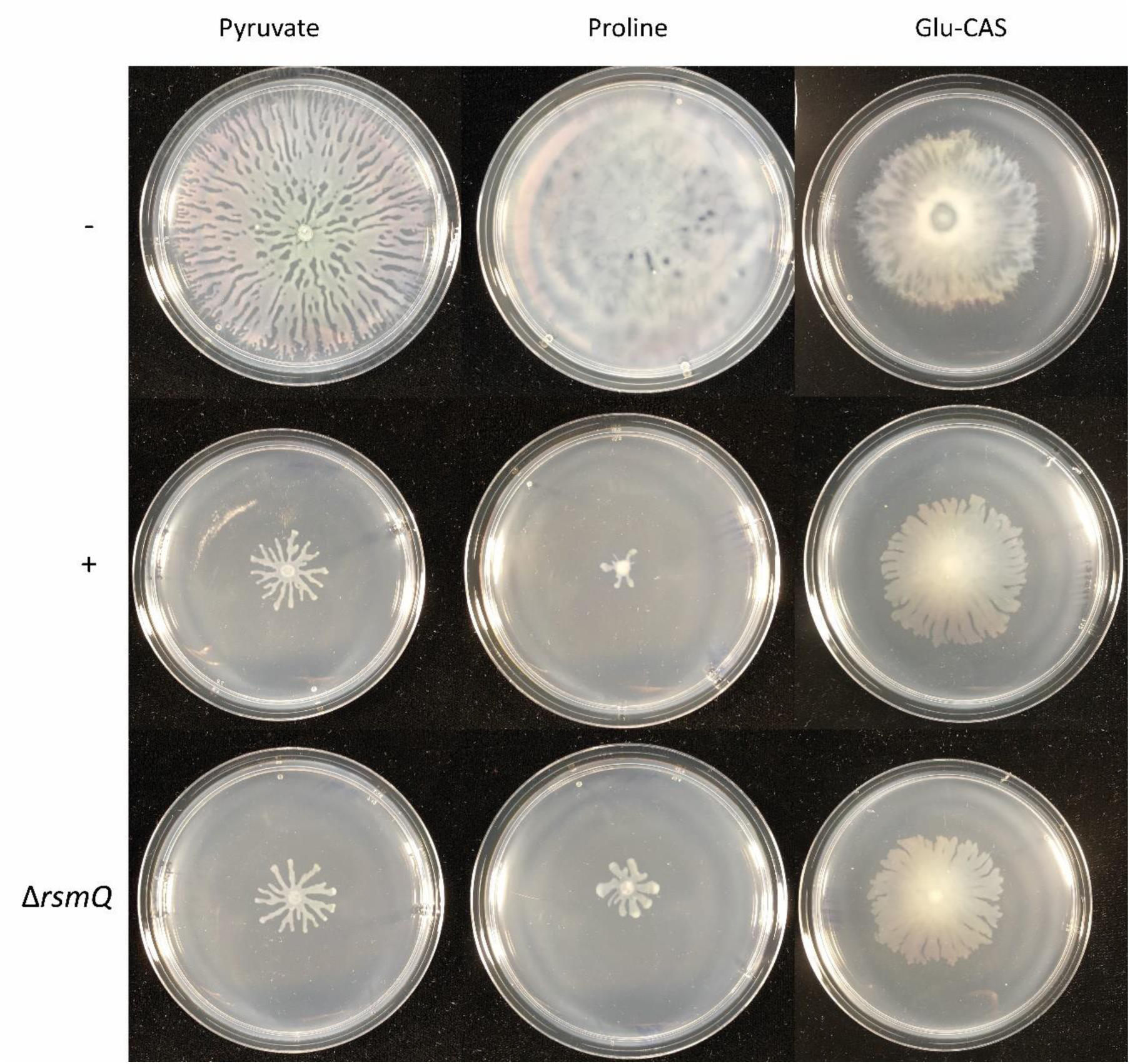
a) Swarming motility after 72 h for SBW25 cells either plasmid free (-) or carrying pQBR103^Km^ (+) or pQBR103^Km^-Δ*rsmQ* grown on 0.5% M9 media with the carbon source indicated.

**Figure S4:**
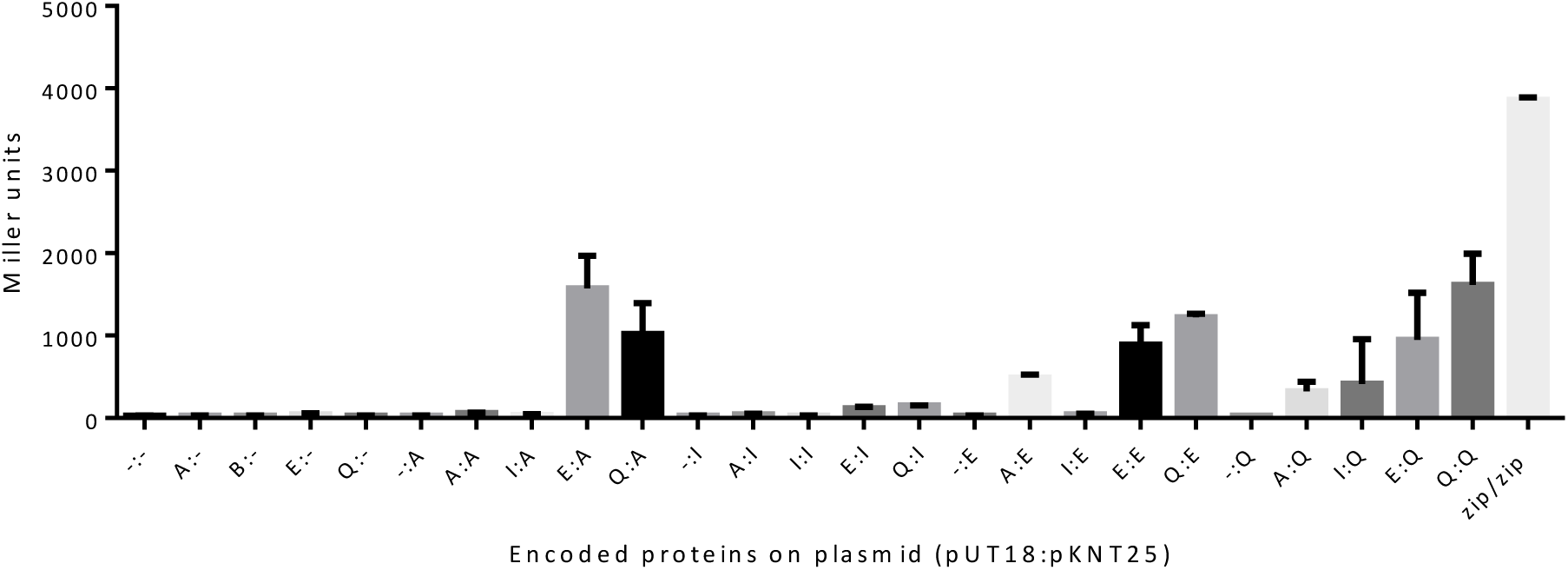
β-galactosidase assay (as shown in figure 5) with all controls shown.

Table S1. Up and Downregulated proteins in for pQBR103^Km^-Δ*rsmQ* / pQBR103^Km^

Uploaded as a separate Excel File (Table S1)

Table S2. BioLog results for SBW25 plasmid free, SBW25 pQBR103^Km^ and pQBR103^Km^-Δ*rsmQ* after 24 hours of growth. Absorbance measured at 590 nm.

Uploaded as a separate Excel File (Table S2)

**Table S3.**
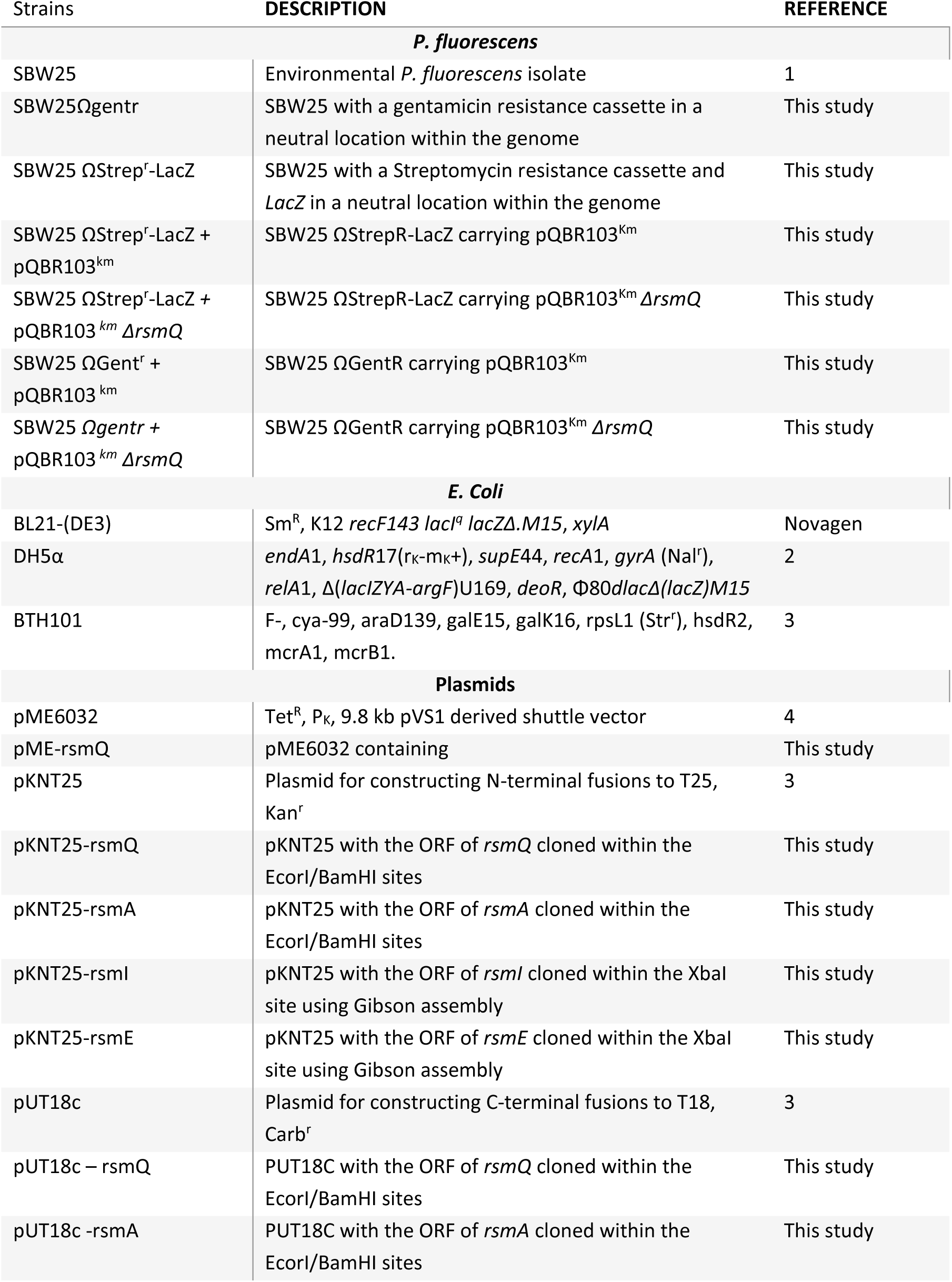
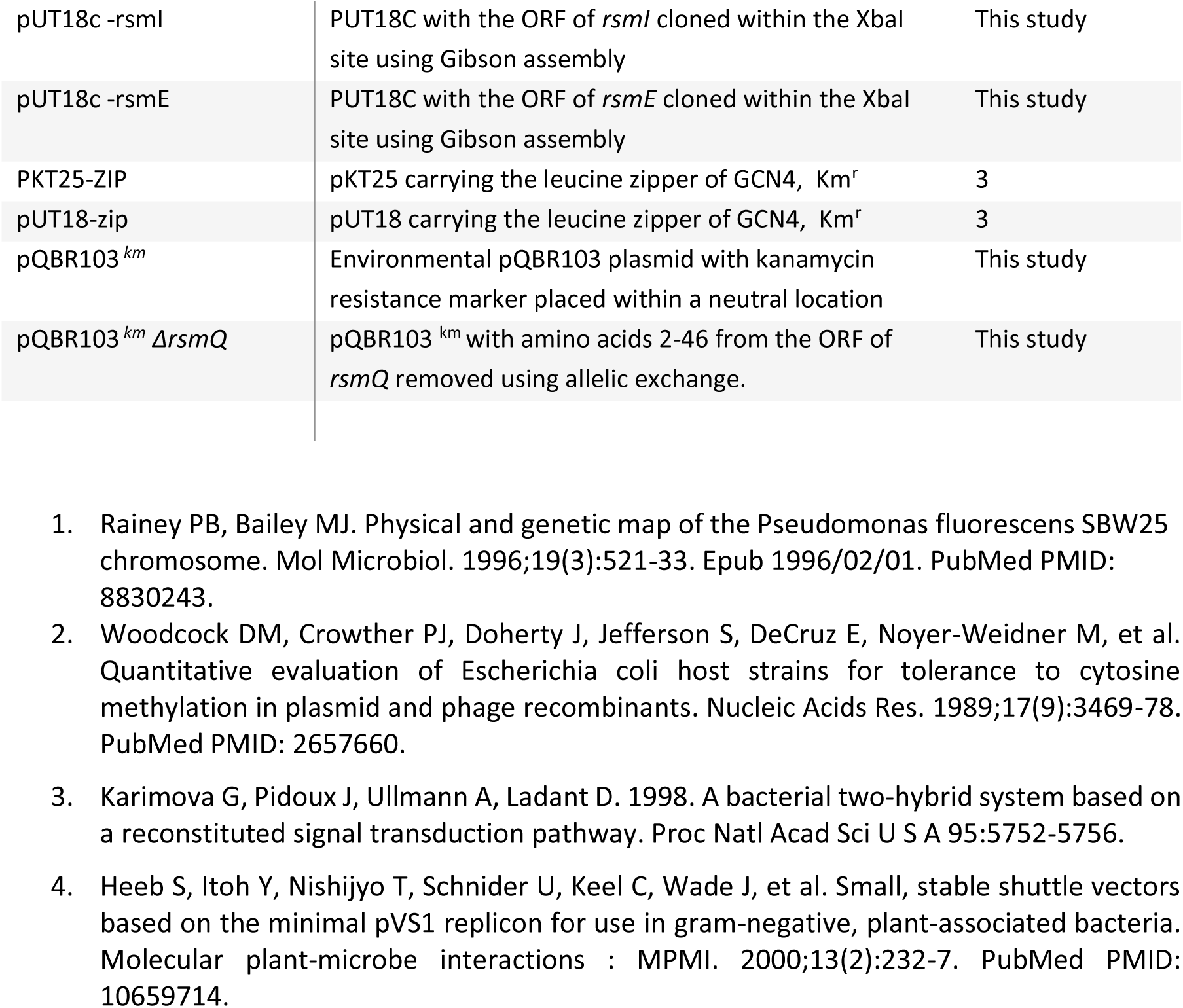
Table of Strains and Plasmids

Table S4. Table of Primers

Uploaded as a separate Excel File (Table S4)

